# A general-purpose protein design framework based on mining sequence-structure relationships in known protein structures

**DOI:** 10.1101/431635

**Authors:** Jianfu Zhou, Alexandra E. Panaitiu, Gevorg Grigoryan

## Abstract

The ability to routinely design functional proteins, in a targeted manner, would have enormous implications for biomedical research and therapeutic development. Computational protein design (CPD) offers the potential to fulfill this need, and though recent years have brought considerable progress in the field, major limitations remain. Current state-of-the-art approaches to CPD aim to capture the determinants of structure from physical principles. While this has led to many successful designs, it does have strong limitations associated with inaccuracies in physical modeling, such that a robust general solution to CPD has yet to be found. Here we propose a fundamentally novel design framework—one based on identifying and applying patterns of sequence-structure compatibility found in known proteins, rather than approximating them from models of inter-atomic interactions. Specifically, we systematically decompose the target structure to be designed into structural building blocks we call TERMs (tertiary motifs) and use rapid structure search against the Protein Data Bank (PDB) to identify sequence patterns associated with each TERM from known protein structures that contain it. These results are then combined to produce a sequence-level pseudo-energy model that can score any sequence for compatibility with the target structure. This model can then be used to extract the optimal-scoring sequence via combinatorial optimization or otherwise sample the sequence space predicted to be well compatible with folding to the target. Here we carry out extensive computational analyses, showing that our method, which we dub dTERMen (design with TERM energies): 1) produces native-like sequences given native crystallographic or NMR backbones, 2) produces sequence-structure compatibility scores that correlate with thermodynamic stability, and 3) is able to predict experimental success of designed sequences generated with other methods, and 4) designs sequences that are found to fold to the desired target by structure prediction more frequently than sequences designed with an atomistic method. As an experimental validation of dTERMen, we perform a total surface redesign of Red Fluorescent Protein mCherry, marking a total of 64 residues as variable. The single sequence identified as optimal by dTERMen harbors 48 mutations relative to mCherry, but nevertheless folds, is monomeric in solution, exhibits similar stability to chemical denaturation as mCherry, and even preserves the fluorescence property. Our results strongly argue that the PDB is now sufficiently large to enable proteins to be designed by using only examples of structural motifs from unrelated proteins. This is highly significant, given that the structural database will only continue to grow, and signals the possibility of a whole host of novel data-driven CPD methods. Because such methods are likely to have orthogonal strengths relative to existing techniques, they could represent an important step towards removing remaining barriers to robust CPD.

## 1 Introduction

Proteins orchestrate molecular life. For this reason, the robust engineering of this class of macromolecules is highly desirable from a multitude of perspectives. It would advance the development of protein therapeutics, it would enable facile design of reagents for biological research, and it could even provide inroads towards new classes of materials. Computational protein design (CPD) could be a particularly attractive means of fulfilling the need for such robust engineering, but CPD techniques have thus far lacked the reliability needed to incorporate them as “black-box” tools in downstream research and technology development. The basic idea behind the modern approach to CPD is to capture structural phenomena (e.g., folding and binding) from physical principles. Since the initial demonstration of this concept by the Mayo group in the late 1990’s (1), many groups have implemented significant advancements on the idea. Notably, the Baker lab developed and has continually refined the widely used Rosetta modeling suite, forming an entire community of researchers and programmers actively contributing to the project (2, 3). Other modern CPD implementations include OSPREY by the Donald lab (4), ORBIT by the Mayo lab (5, 6), PROTEUS by Simonson and co-workers (7), and FoldX by Serrano and colleagues (8). Advancements introduced over the years by these and other researchers have aimed to improve the treatment of a range of physical effects towards a more realistic representation of proteins, including modeling backbone flexibility (9), better accounting for side-chain degrees of freedom (10, 11), treatment of bulk-solvation and individual-water effects (12–14), consideration of ensemble properties of structure (15–18), scoring function improvements (3, 19), modeling the unfolded state (15, 20, 21), and others. Several recent reviews provide excellent summaries of achievements and challenges in CPD, including by Donald and co-workers (22) (a thorough analysis of algorithmic aspects), by Ilan Samish and Regan *et al*. (23, 24) (placing current work in a historical perspective), and Woolfson *et al*. (25) (a commentary on the implications of the ability to design novel structures). Nevertheless, despite all of these developments, and many examples of successful designs in the literature (18, 26–35), it is still the case that CPD methods are not robust. State-of-theart techniques, even in the hands of experts, fail too frequently, showing that significant inaccuracies are still present in the underlying models and motivating the development of alternative solutions.

At first glance, the reason for this lack of robustness is the stiff tradeoff between physical accuracy and computational feasibility. That is, even if we assume that standard molecular mechanics (MM) provides enough accuracy to capture basic protein thermodynamics, the full consideration of MM (and the implied statistical mechanics) is computationally infeasible in the context of CPD. Thus, CPD models must necessarily admit severe simplifying approximations and are inaccurate. While this logic is certainly true, it is worth considering that an even deeper reason behind the low robustness of modern CPD may be inherent in attempting to synthesize relationships between sequence and structure from models of interatomic interactions. Whereas the latter undoubtedly defines the former, the two are many steps removed, making it difficult in practice to tune models (i.e., atomistic interaction parameters) to achieve the correct sequence-structure mapping across all systems. And this is even before we consider that MM may not, in fact, be capable of capturing all relevant sequence-structure effects in a single model (36).

In this work we consider the possibility of performing protein design by directly observing and learning from sequence-structure relationships present in available protein structures, rather than aiming to synthesize them from atomistic interactions. This type of methodology is likely to have entirely orthogonal strengths and weaknesses, relative to the standard CPD approach. If sufficient structure and sequence data are available, a data-driven approach may be difficult to outperform in terms of robustness. But it is unclear what “sufficient” means in this context and how close we may be to this threshold today. Thus, the two main objectives of this work are: 1) to develop a general-purpose CPD framework that relies solely on previously available protein structures, and 2) thoroughly benchmark this framework as a means of understanding to what extent the present Protein Data Bank (PDB) is sufficiently large to support general-purpose practical protein design.

As of August 2018, over 140,000 entries have been deposited into the PDB, with a yearly increase of ∼10,000. Experimental structures have always been a key source of fundamental insights on protein structure-sequence relationships, with degeneracies in structure space—i.e., repeated structural patterns or motifs and associated sequence preferences—proving especially insightful. Examples of such degeneracies include basins of contiguous backbone conformation with their sequence propensities (37, 38), side-chain rotamers and their dependence on backbone dihedral angles (11), or loop geometries with associated sequence preferences (39–41). In a recent study, we showed that structural degeneracy extends beyond local-in-sequence motifs (e.g., backbone fragments) or simple geometric descriptors (e.g., distances, dihedral angles, or degrees of burial) and into tertiary and quaternary structural geometries (42). Specifically, we found that local-in-space motifs, which we dubbed TERMs (<u>ter</u>tiary <u>m</u>otif<u>s</u>), are highly recurrent in the structural universe. For example, we found that only ∼600 TERMs are sufficient to describe over 50% of the observable structural universe, and that TERMs tend to repeat many times, across proteins apparently unrelated by homology (42).

Here we present a CPD framework dTERMen (<u>d</u>esign with <u>TERM</u> <u>en</u>ergies) that takes advantage of this degeneracy. It systematically breaks the target structure into its constituent TERMs and accounts for the sequence preferences of each by analyzing sequences of closely matching backbone fragments identified from the PDB using our structure search engine MASTER (43). With this information, a sequence-level pseudo-energy table is generated, enabling the scoring of any sequence for compatibility with the target backbone, identification of the optimal sequence, and other optimization or search tasks. We previously showed that PDB-derived TERM-based sequence statistics are able to distinguish correct from incorrect structural models and predict the impact of amino-acid substitutions on protein stability on par with or better than state-of-theart methods built for these tasks (44, 45). Thus, there are reasons to believe that a TERM-based design framework may be feasible. Here we aim to establish this definitively.

The concept of data-driven CPD has been explored before. First, any statistical potential can be placed into this category of techniques, such that essentially any existing CPD method can be thought of as partially data-driven, as the use of statistical pseudo-energies is widespread (e.g., rotamer probabilities, backbone phi/psi propensities, residue/atom-pair interaction pseudo-energies, and so on). ABACUS is an example of a fully statistical energy function for fixed-backbone protein design (46–48). This rotamer-level statistical energy function includes up to two-body pseudo-energies, computed from probabilities of observing rotamers (or rotamer pairs) in the PDB conditioned on structural properties of the position(s) involved being similar to the structural properties of the target residues. These properties include backbone torsional angles, solvent accessibility, and inter-positional backbone geometry (in the case of pair energies). The method has been tested both *in silico* as well as by designing and experimentally validating a novel sequence intended to fold to the structure of human ubiquitin (46).

PANENERGY is an example of atomistic statistical energy function built for CPD and related applications (49). It builds upon a traditional atom-pair PDB-derived statistical interaction energy, but applies a novel “connectivity scaling factor” to modulate inter-atomic interactions. The factor is based on the degree of chemical connectivity of interacting atoms, which effectively emulates course-graining (thus introducing some many-body character to the underlying atomic pairwise interaction model) without loss of atomistic detail. The function is further supplemented with an amino-acid environment statistical energy and a non-polar surface area-based cavitation term (49). PANENERGY has been shown to perform well on predicting mutational protein stability changes and protein-ligand binding free energies, decoy discrimination, and native sequence recovery (49).

Zhang and co-workers have developed an evolution-based statistical CPD approach, called EvoDesign (50–52). The method works by identifying structural analogs of the target scaffold from the PDB library, via TM-align (53), and deriving a multiple sequence alignment from these. A position-specific scoring matrix (i.e., structural profile) is then generated according to the extracted evolutionary relationships. This structural profile is used to generate proposed sequences, which are then scored based on predicted secondary structure, backbone torsion angles, and solvent accessibility. EvoDesign has been tested computationally and several designed proteins have been shown to have circular dichroism and NMR spectra consistent with well-ordered structures.

Keating and co-workers designed coiled-coil peptides that selectively bind to basic-region leucine-zipper (bZIP) domains of human proteins using a scoring function generated from a large body of experimental coiled-coil dissociation constants via machine learning (54). Guided by this function, novel coiled-coil peptides were assembled from short heptad sequence fragments derived from natural bZIP proteins. More recently, Qi and coworkers applied deep-learning neural networks to CPD by expressing it as the problem of predicting the amino-acid identity at a position given its structural neighborhood (55). Specifically, the neural network comprised an input layer, several hidden layers, and an output softmax layer, with basic geometric and structural properties (e.g., Cα–Cα distance, secondary structure, and solvent accessible surface area) fed as input features. The output of the softmax layer were the probabilities of the 20 natural amino acids at the analyzed target position. The neural network was trained with the aim of “predicting” the native amino acid as frequently as possible, exhibiting an accuracy of 38% in a five-fold cross validation.

A fundamental difference between dTERMen and prior data-driven approaches lies in the underlying assumption as to how sequence is dictated by structure—i.e., the TERM hypothesis. This assumption derives from prior observations of the “quantized” or “digital” nature of protein structure, at the local (in space) level (42). Thus, dTERMen differs from statistical potentials in that it goes beyond simple geometric descriptors and analyzes apparent sequence preferences in the context of larger well-defined backbone motifs, relying on their quasi-digital nature. On the other hand, it is different from machine-learning (ML) approaches in that the TERM hypothesis effectively serves as a strong “prior” on the functional form of the model, which allows the method to bridge data sparsity issues. Further, unlike ML-based methods, the goal of dTERMen is not to predict native-like amino acids from a set of structure-defining inputs. Rather, the method seeks a residue-level effective potential that maximally bears out the TERM hypothesis.

These differences afford dTERMen several advantageous features. In contrast to ML, it does not necessitate a model training step. The modularity of TERMs enables dTERMen to exploit entirely unrelated protein structures, broadening its applicability to a great extent. Further, the potential for increased accuracy is effectively built into the method. More structures in the database produce more accurate and refined sequence preferences and, ultimately, more accurate sequence landscapes. Thus, we can expect better performance with time, as the PDB continues to grow.

In this study, we undertake a thorough validation of dTERMen. We demonstrate that dTERMen produces native-like sequences from native backbones and that dTERMen-designed sequences are predicted to form the intended structure nearly as often as native sequences. We also show that dTERMen pseudo energies correlate with experimental stability metrics and predict design success. Further, we validate dTERMen experimentally by designing an entirely novel surface for Red Fluorescent Protein mCherry. The single tested variant, whose sequence was recognized as the optimal by dTERMen, folded, was monomeric, showed similar chemical stability as mCherry, and even exhibited fluorescence. These positive results certainly validate the specific computational framework proposed in this work. More importantly, however, they validate the general notion that structural data accumulated in the PDB are now sufficient to enable general-purpose CPD based on mining sequence-structure relationships in the contexts of motifs extracted from the target protein. This is a fact of fundamental importance for the field, in our opinion, as it signals the possibility of an entirely new set of protein design and modeling approaches that will be largely orthogonal to prior developments, with complementary strengths and weaknesses. This should have a positive effect on advancing the issue of low robustness of CPD.

## 2 Results

Results are organized as follows: section 2.1 summarizes the philosophy behind dTERMen (full details in Materials and Methods), sections 2.2 – 2.6 describe a series of computational benchmarks of the method, and section 2.7 presents the results of applying dTERMen to the total surface redesign of mCherry. Details of all experimental and computational procedures are provided in Materials and Methods.

### 2.1 dTERMen procedure

Suppose we are given a target protein structure, *D*, for which we need to find an appropriate amino-acid sequence. The goal of our procedure is to define effective self energies for each amino acid at each position of *D* and effective pair (or higher-order) interaction energies between amino acids at pairs of positions (or larger clusters). We collectively refer to these as energy parameters (EPs) and their values are deduced from statistics of structural matches to appropriately defined TERMs comprising *D*. The matches are obtained by searching within a redundancy-pruned subset of the Protein Data Bank (PDB), although more specialized databases can also be employed (e.g., when designing transmembrane proteins, one may want to restrict the database to only such proteins).

The procedure recognizes several types of effective energetic contributions at play in defining protein sequence-structure relationships: the propensity of an amino-acid residue for the general environment of a position, such as the burial state (environmental energy); interactions between an amino acid at a position and its surrounding backbone, which are further broken into contribution from its local-in-sequence backbone fragment (the own-backbone component) and contributions from spatially-proximal backbone fragments (the near-backbone component); and interactions between pairs and higher-order clusters of amino acids. Environment, own-, and near-backbone energies are self contributions, whereas the remaining ones constitute pair and higher-order contributions.

Once *D* is appropriately decomposed into a set of overlapping TERMs (see below), and structural matches are identified for each TERM from the database, EP values are deduced following two general principles. **Principle 1** states that sequence statistics within TERM matches are driven only by the EPs involving positions contained in the TERM (e.g., a pair EP influences the statistics of a TERM if and only if the corresponding pair of positions are contained within the TERM). This assumption is reasonable in cases where the matches arise from a large diversity of structural backgrounds, such that context effects average out. Certain redundancy-removal steps are key to making sure that this assumption holds well in practice (see Materials and Methods). It follows from principle 1 that EP values should be sought to maximally describe the sequence data observed in TERM matches. **Principle 2** stipulates that higher-order parameters be involved only when needed—i.e., models involving only lower-order parameters are preferred, all else being equal. This means that higher-order EPs act as correctors to lower-order contributions. For example, pair energies are needed only to describe those aspects of sequence statistics that are not satisfactorily described with self contributions.

#### 2.1.1 TERM decomposition: main idea

Within the sequence space compatible with folding into *D*, some residue pairs are *coupled*—i.e., the optimal amino-acid identity of one residue depends on the identity of the other. Such coupled positions can be identified through the structure of *D*, by finding position pairs capable of hosting amino acids that have an influence on each other via direct or indirect physical interactions (see Materials and Methods). In addition, in some systems with sufficiently large multiple sequence alignments, evolutionary co-variation can suggest coupled positions. Finally, experimental evidence identifying specific coupled position pairs may also be available.

Whatever the source of the inference, the coupling relationships in *D* can be thought of as an undirected graph, where nodes represent residues and edges signify coupling, with edge weights optionally indicating the strength of coupling (inferred from structure or known); let us call this graph *G*. The final pseudo-energy model should involve self contributions for all nodes, pair contributions for all edges, and (optionally) higher-order contributions for a subset of connected sub-graphs of *G*. Further, to describe near-backbone interactions, we define a directed graph, *B*, in which nodes represent residues and a directed edge between *a* and *b, a* → *b*, signifies that the backbone of *b* can influence the amino-acid choice at *a*. As with coupling, such pairs of positions can be identified through a structural analysis of *D* (see Materials and Methods). A TERM decomposition of *D* should respect the structures of *G* and *B* and enable the extraction of EPs for the above contribution types. Specifically, a complete set of TERMs describing *D* must be such that every residue and every pair of coupled residues be covered by at least one TERM. In addition, if higher-order coupling contributions are desired, TERMs covering corresponding connected sub-graphs of *G* should be included as well. Similarly, if higher-order near-backbone contributions for position *i* are desired, TERMs covering *i* and all (or a subset of) nodes to which it has directed edges in *B* should be included as well.

#### 2.1.2 TERM decomposition: specifics

Here we describe the specific TERM decomposition procedure used in this study (see Fig. 1), noting that many other procedures that follow the above principles can be appropriate. We define TERMs via connected sub-graphs of *G* or *B*. If a subgraph includes the node corresponding to residue *i*, then the resulting TERM includes residues (*i* − *n*) through (*i* + *n*), where *n* is a parameter (we generally use *n* = 1 or 2, and exclusively *n* = 1 in this study); see Fig. 1. We first define a TERM for each node in isolation (i.e., treating it as a one-node subgraph); we refer to these as singleton TERMs. Singletons are used to deduce own-backbone contributions (see below). Next, to capture near-backbone contributions at residue *i*, we create a TERM that involves node *i* and all nodes to which it has directed edges in *B*; lets call this set *β*(*i*)—the “influencing” residues. If such a TERM does not have a sufficient number of close structural matches in the database (see Materials and Methods for details of match definition), the effect of the neighboring backbone on *i* needs to be captured with multiple TERMs. In particular, we start by defining TERMs containing *i* and each residue in *β*(*i*), independently. Of these TERMs, the one with the most structural matches (suppose it is the one containing nodes *i* and *j* ∈ *β*(*i*)) is chosen for expansion, with each remaining node *k* ∈ *β*(*i*) ∩ *j* considered for inclusion into the sub-graph, one at a time. Once again, of these, the one with the most matches is selected, and this procedure is repeated until no more nodes can be included into the growing sub-graph. Once this occurs, the expanded TERM is accepted into the overall TERM decomposition, with all of the influencing residues involved in it marked as covered. The procedure is then repeated, using only uncovered influencing residues, until all residues in *β*(*i*) are covered. This technique is a generalization of considering a single TERM that covers *i* and all *β*(*i*), splitting the near-backbone effect into as few TERMs as needed to retain sufficiently good sequence statistics, while capturing as much of the near backbone environment simultaneously as possible. TERMs generated for capturing near-backbone effects are referred to as near-backbone TERMs (see Fig. 1).

**Fig. 1.**
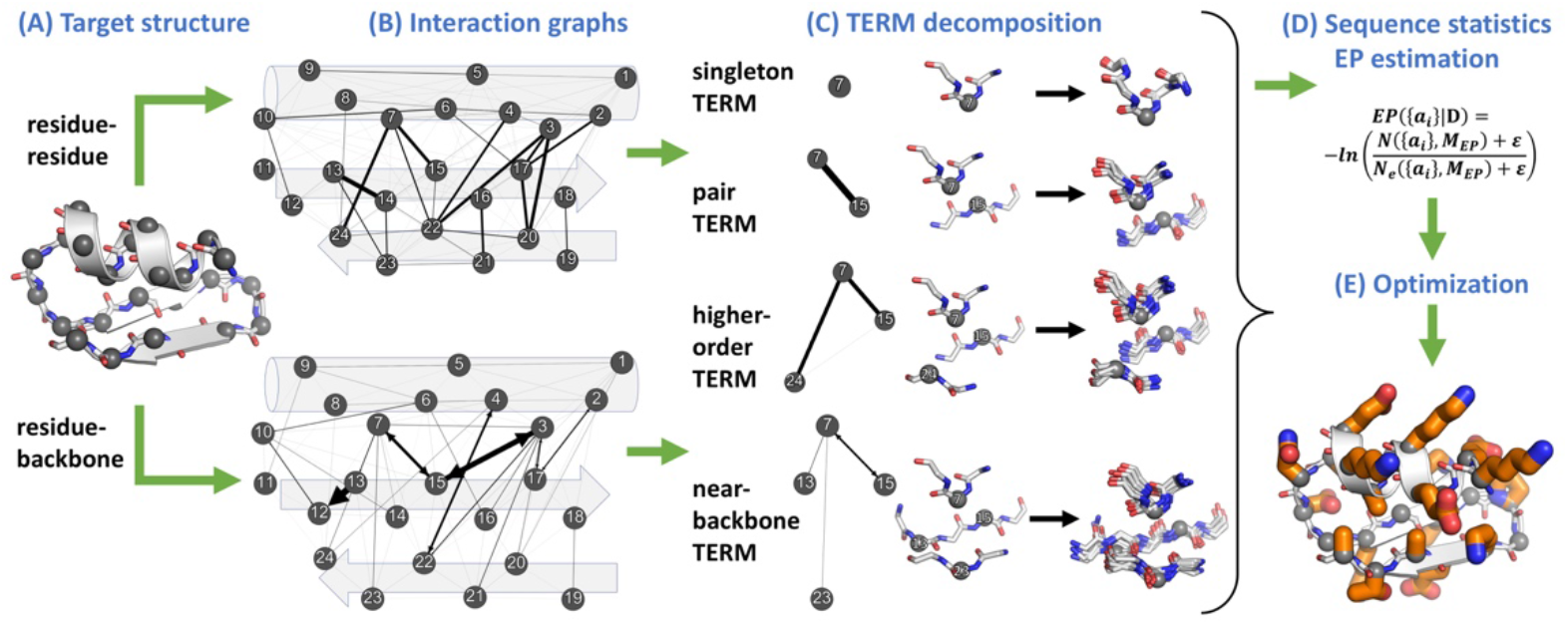
Diagram of dTERMen procedure. Target structure (A) is decomposed into TERMs guided by the graph of its coupled residues, (B) top, and the graph of residue-backbone influences, (B) bottom. Close matches to each TERM from the structural database are identified, (C), and the sequence alignments implied by these matches are used to estimate EPs governing the sequence-structure relationship in the target structure, (D). Combinatorial optimization is then used to produce the optimal sequence for the target, (E), or can also be used to build a library of design variants or for other tasks.

We next define one TERM for each pair of nodes in *G* connected by an edge. These are referred to as pair TERMs and used to deduce pair interaction EPs. Finally, higher-order TERMs are defined for select connected sub-graphs of *G* and used in deducing higher-order interactions. Individual higher-order sub-graphs can either be chosen manually, based on prior knowledge of the system or inspection of structure *D*, or automatically using an appropriate structure-based rule (e.g., only fully connected three-residue sub-graphs, potentially filtered by edge weights). These TERMs are only included if they possess a sufficient number of structural matches (see Materials and Methods for details). While our method can extract higher-order couplings (provided enough data are available), we have generally found it unnecessary to do so in practice, and all of the examples presented in this study included only up to pair contributions.

#### 2.1.3 Computing EPs

Following the two general principles outlined in section 2.1, many specific computational procedures can be formulated to extract EP values from the data provided by a TERM decomposition (i.e., TERM matches and their sequence statistics). Here we employ a procedure that considers pseudo-energetic contributions in a hierarchy, with each next type of contribution introduced only to describe what is not already captured by previous ones. By including higher-order contributions later in the hierarchy, we make sure that these are only used as correctors (to the extent necessary) over what is already described by lower-order contributions. Further, the earliest contributions in the hierarchy are those associated with the strongest sequence statistics, such that highest-confidence effects are captured first, relatively unaffected by statistical noise. The specific order of contributions in the hierarchy used here is: 1) amino-acid backbone *φ/ψ* dihedral angle propensities, 2) amino-acid backbone *ω* dihedral angle propensities, 3) pseudo-energy associated with the general environment (burial state) of a residue, 4) own-backbone contributions, 5) near-backbone contributions, 6) residue pair contributions, and 7) any considered higher-order contributions. The details of pseudo-energy calculation are presented in Materials and Methods.

### 2.2 dTERMen predicts native-like sequence from NMR or X-ray backbones

“Native sequence recovery” is a classical benchmark experiment for protein design methods (21). In this experiment, the procedure in question is applied to design sequences for backbones taken from experimentally determined native-protein structures. The idea is that a good method should propose sequences that are similar to the corresponding native sequence. Indeed, the native sequence is one solution that is known to fold to the targeted conformation. And whereas many other sequences likely can do so as well, it is reasonable to assume that there ought to be some similarity between any sequence that folds to the specified backbone and the native sequence.

To measure native sequence recovery, we curated a set of 90 X-ray structures and 31 NMR structures of globular proteins, ranging in length from 50 to 150 residues (see Materials and Methods). dTERMen was applied to design a sequence for each backbone, and the globally optimal sequence obtained by Integer Linear Programing (ILP) optimization was compared with the native sequence. For comparison, the same backbones were also used in designs by Rosetta, using the *talaris2013* energy function (56) (see Materials and Methods). Table I summarizes the resulting native sequence recovery rates. The two methods perform similarly, with dTERMen giving slightly less native-like sequences for X-ray backbones (∼29% relative to Rosetta’s ∼33%, on average) and slightly more native-like ones for NMR backbones (∼24% relative to Rosetta’s ∼22%, on average). Thus, dTERMen performs on par with the state of the art, of which Rosetta Design is a great representative. Interestingly, however, the specific sequences proposed by dTERMen and Rosetta are quite different (see last two rows of Table I). This is in line with the fact that the two methods choose sequences based on entirely different principles, but it makes the comparable performance on native sequence recovery more interesting. It is noteworthy that the rates exhibited by either method map roughly into the “twilight” zone of sequence identities, where assurance of structural homology is known to drop quickly with sequence similarity (57). This is consistent with the fact that both methods aim to pick a sequence that is merely compatible with folding to the desired backbone, without any additional constraints of functionality or evolutionary history that may shape the specific native sequence.

**Table I.** dTERMen and Rosetta propose distinct, similarly native-like sequences given native backbone. Shown are means and standard deviations of sequence identities between designed and native sequences, within respective datasets. The last two rows show the rate of recovering disulfide bonds (i.e., two cystine residues designed at locations occupied with disulfide-bonded cystines in the native structure).

|  | Sequence identity |  |
| --- | --- | --- |
|  | X-ray (90) | NMR (31) |
| dTERMen vs. native | 28.6% ± 5.8% | 23.9% ± 6.1% |
| Rosetta vs. native | 32.6% ± 6.9% | 22.2% ± 6.3% |
| dTERMen vs. Rosetta | 26.8% ± 5.7% | 22% ± 4.1% |
|  | Disulfide identity |  |
| dTERMen vs. native | 24/80 (30%) | 1/7 (14.3%) |
| Rosetta vs. native | 0/80 (0.0%) | 0/7 (0.0%) |

That dTERMen exhibits somewhat higher native sequence recovery rates on NMR backbones, compared to Rosetta, is consistent with the “fuzzier” interpretation of backbone coordinates by the former (as sequence statistics are discovered in the context of similar, but not identical backbone structural matches). To investigate this apparent insensitivity to backbone noise, we compared sequences designed on alternative NMR backbones as well as those designed on X-ray and NMR structures of the same protein (see Materials and Methods). Alternative NMR models or X-ray vs. NMR structures of the same protein can be seen as conformations that are, to within experimental error, equivalent and equally appropriate for the native sequence. An ideal CPD method should recognize this and design very similar sequences given such backbones. As shown in Table II, dTERMen is quite consistent across such experimentally equivalent backbones, producing sequences with 40–50% sequence identity, on average. Rosetta, on the other hand, shows much greater variability, with sequence identities from equivalent backbones in the range of 20–30%.

**Table II.**
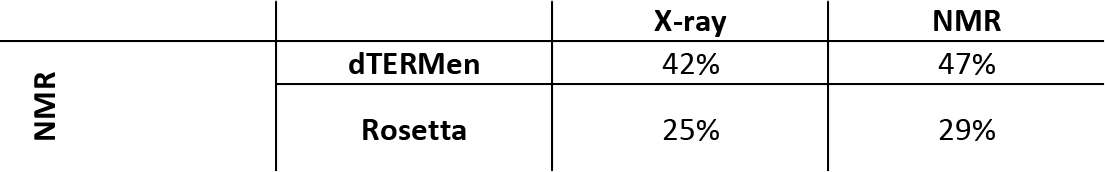
dTERMen is consistent with respect to small backbone adjustments. Rates are calculated over a dataset of 11 matched pairs of NMR and X-ray structures of the same proteins (see Materials and Methods). For NMR-vs-NMR comparisons, all pairs of models for each NMR entry are considered and averaged overall all 11 NMR structures. For NMR-vs-X-ray comparisons, each NMR model is compared to the corresponding X-ray structure, and averages are calculated over all 11 pairings.

A closer look at native sequence recovery based on the degree of burial reveals that Rosetta’s high performance for X-ray backbones is dominated by core positions, where the method achieves the very high rate of ∼52%, on average, whereas the performance of dTERMen is more uniform across position types (see Table SI). The performance of the two methods is comparable for interfacial positions and dTERMen produces slightly higher rates for surface positions (see Materials and Methods for position type definitions). Relative trends are similar for NMR structures, with the overall performance shifted towards dTERMen (Table SI).

As shown in the last two rows of Table I, dTERMen has a high rate of disulfide bond recovery—e.g., 24 out of 80 disulfides (30%) were recovered from X-ray structures (the rate appears lower for NMR structures, but it is out of only seven disulfides occurring in this set). The rate seems especially high when considering that it refers to the simultaneous recovery of two residues (and thus, on average, we would expect it to be ∼8–9% based on the per-position recovery rates in Table I). This indicates that there is significant mutual information between disulfide-bonded positions that dTERMen captures. In contrast, Rosetta did not recover any disulfide bonds; this is not surprising, because: 1) Cys is a rare amino acid, so Rosetta is parameterized to choose it infrequently, 2) simultaneous recovery of two disulfide-bonded cystines is made even more unlikely by the fact that in their bonded conformation the two residues would produce a significant van der Waals clash, and 3) standard scoring functions (such as the *talaris2013* used here) do not include disulfide bond terms. Of course, dTERMen also does not have any special provisions for disulfide bonds, but it chooses these (“unknowingly”) by simply obeying the sequence-structure patterns observed in the database.

### 2.3 dTERMen-designed sequence predicted to fold to desired structures

The ultimate goal of a design method is to propose sequences that fold into the desired structure, and not to reproduce the most native-like sequences given native backbones. Folding into the correct structure involves not only forming favorable interactions in the context of the target backbone, but also requires the sequence to disfavor the multitude of available alternative conformations. This last property, which has been referred to as “fold specificity” (58), is particularly difficult to achieve in CPD and this is likely a key reason behind many design failures. The best way to assess this and other qualities of a designed sequence is to characterize it experimentally (e.g., see section 2.6), but this is costly and time consuming. Another way is to assess whether designed sequences are predicted to fold into the desired structure *in silico*, using a cutting-edge structure prediction method. Though this is not nearly as costly as experiments, it does not provide ground truth. *De-novo* structure prediction is a difficult problem, with a high rate of errors. So, the mere prediction that a designed sequence either does or does not fold into the desired structure cannot be given significant weight. On the other hand, if such a test-by-structure-prediction is performed on a large set of designed sequences, emerging from diverse templates, and used to compare two methods, then statistically significant differences in performance can be interpreted as meaningful. This is because the two methods would act as controls for one another, with the assessment of either subject to the same difficulties and errors associated with structure prediction.

Using the designed sequences targeting the native backbones from the previous section, we performed *de-novo* structure prediction with each using a standalone copy of I-TASSER, making sure that data from homologues of the protein whose backbone was used as the target did not contribute to the calculation (see Materials and Methods). Each I-TASSER run, which took ∼20 CPU hours on average, was asked to produce 10 models and each was subsequently compared with the desired target structure to extract its TM score (59). Each dTERMen and Rosetta designed sequence was subjected to the same treatment, with Fig. 2A comparing the results. As expected, TM scores were not usually close to 1.0, which represents both the difficulty of structure prediction and the fact that some designs may not fold into the desired structure. However, dTERMen design performed better, on average, with their TM scores exceeding the TM-score of the corresponding Rosetta design in 58% of cases. The mean TM-scores over dTERMen and Rosetta designs were 0.48 and 0.45, respectively (p = 0.003), with medians showing a similar trend (see Table III). Furthermore, 43.2% of dTERMen designs exhibited a TM score over 0.5 (a value typically chosen for delineating a roughly correct fold), and only 38% of Rosetta designs reached this value. Models derived from dTERMen sequences also exhibited higher fractions of correct secondary structure types (see Fig. 2B).

**Fig. 2.**
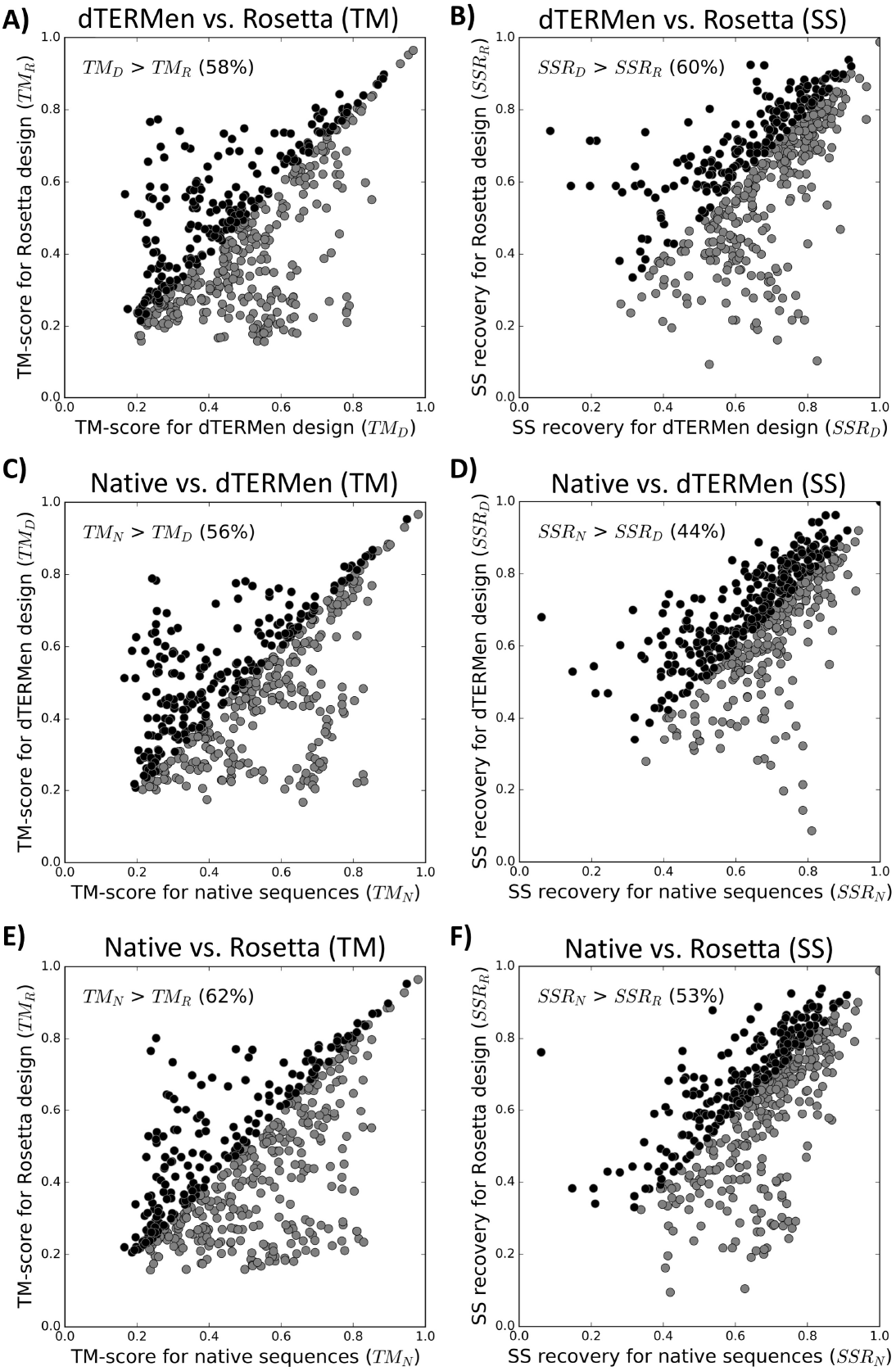
Testing of dTERMen-designed sequences in structure prediction using I-TASSER. Structures were predicted for three sequences corresponding to each target structure (dTERMen-designed, Rosetta-designed, and native), with I-TASSER being asked to predict top 10 models. Models for each sequence were numbered (in the order returned by I-TASSER), allowing us to compare the *i*-th model between any two sequences (e.g., the top model by dTERMen versus Rosetta). Each point in each plot represents a comparison between some model *i* (*i* ∈ [1; 10]) for two sequences from the same template (gray and black points map below and above the diagonal, respectively). **A)** and **B)** compare dTERMen and Rosetta sequences, **C)** and **D)** compare native and dTERMen sequences, and **E)** and **F)** compare native and Rosetta sequences. In **A), C)**, and **E)**, the comparison is by TM-score of the model relative to the native structure; in **B), D)**, and **F)**, the comparison is by fraction of residues with the correct secondary-structure classification. The legend of each plot indicates the fraction of times one set of compared sequences outperforms the other.

**Table III.** Summary of structure-prediction performance of dTERMen-designed, Rosetta-designed, and native sequences. 1—fraction of models built from either sequence set that achieved a TM-score above 0.5 (relative to the native structure). 2— mean TM score across all predicted models within each sequence set. The p-values for the null hypothesis that the true means of underlying distributions are identical are 0.05 for comparing dTERMen and native sequences, 0.003 for comparing dTERMen and Rosetta sequences, and 0.000002 for comparing Rosetta and native sequences. 3— median TM score across all predicted models within each sequence set.

|  | % with TM > 0.5<sup>1</sup> | Mean TM<sup>2</sup> | Median TM<sup>3</sup> |
| --- | --- | --- | --- |
| Native | 50.7% | 0.508 | 0.503 |
| dTERMen | 43.2% | 0.484 | 0.474 |
| Rosetta | 38.0% | 0.449 | 0.427 |

The above metrics rate dTERMen designs, on average, better than Rosetta ones. But several questions remain. First, how significant is this difference (beyond mere statistical significance)? Second, how good is this performance in an absolute sense? And finally, could it be that our test merely quantifies the “predictability” of sequences by the specific structure-prediction method used that does not correlate with design quality? To help address all of these questions, we ran a control calculation, where all of the above analysis was repeated for native sequences. Because native sequences do, in fact, fold to the desired structure, their performance in the test can be thought of as that of a “perfect” design method, allowing us to quantify both how far from ideal our methods are and how significant their performance differences are. Finally, if our structure-prediction test is not capturing design sequence quality, then we would not expect native sequences to do any better than artificially designed ones.

Figs. 2C,E compare the performance of native sequences with that of dTERMen designs and Rosetta designs, respectively, with summary metrics shown in Table III. Native sequences perform better than both dTERMen and Rosetta, which validates our test, dTERMen is second best, and Rosetta is last. Further, the performance of dTERMen, by all metrics, is about half way between native sequences and Rosetta. For example, 51% of models from native sequences have a TM-score above 0.5, while this number is 43% and 38% for dTERMen and Rosetta sequences, respectively. This suggests that the difference between dTERMen and Rosetta sequences is indeed significant. Finally, the difference between dTERMen and native sequences is at the edge of statistical significance. For example, mean TM-score is 0.51 for native sequences and 0.48 for dTERMen sequences, with the corresponding p-value of 0.05 (see Table III). In fact, in terms of recovery of the correct secondary structures, dTERMen sequences perform slightly better than native ones, while Rosetta sequences perform worse than native ones (compare panels D and F in Fig. 2).

### 2.4 dTERMen imposes native-like amino-acid usage

Good performance in native sequence recovery or structure prediction tests certainly suggests that dTERMen designs are generally appropriate for the targeted backbone. However, a design method can still possess many inappropriate biases even if it performs well in these tests. For example, the method may be particularly poor at choosing surface residues. This may not significantly affect the overall sequence recovery rate (as surface positions are generally recovered poorly) and structure prediction methods may not be sensitive enough to detect poorly designed surfaces (e.g., merely having polar/charged residues could be sufficient). Or, the method may perform poorly with respect to rare amino acids (e.g., it may never choose Cys), again potentially without a substantial penalty on sequence recovery or structure prediction performance.

Thus, as an alternative (and somewhat orthogonal) quality test of designed sequences obtained by dTERMen, we looked at the amino-acid utilization implied by the method, and whether it agrees with the native amino-acid utilization. To this end, we randomly chose 10,000 non-redundant protein structures from the PDB and redesigned one randomly-chosen position in each, keeping the remaining amino acids at their wild-type identity. From this, we estimated conditional probabilities of the form *p*(*X*|*Y*)—the probability of proposing amino acid *X* at a position that is natively occupied by amino acid *Y*. These can be thought of as mutational probabilities, and organized into a 20×20 matrix would form a mutational probability matrix *M*; shown in Fig. 3A. This matrix represents a linear operation that describes how the distribution of amino acids would change under the guidance of dTERMen. We can, therefore, ask what the amino-acid distribution would have to be so as not to be perturbed by *M*; this is just the eigenvector of *M* with the eigenvalue of 1.0. This stationary amino-acid distribution is plotted in Fig. 3C against the native amino-acid distribution found in the PDB. There is a clear and statistically-significant correlation between the two, indicating that dTERMen predicts native-like utilization of amino acids.

**Fig. 3.**
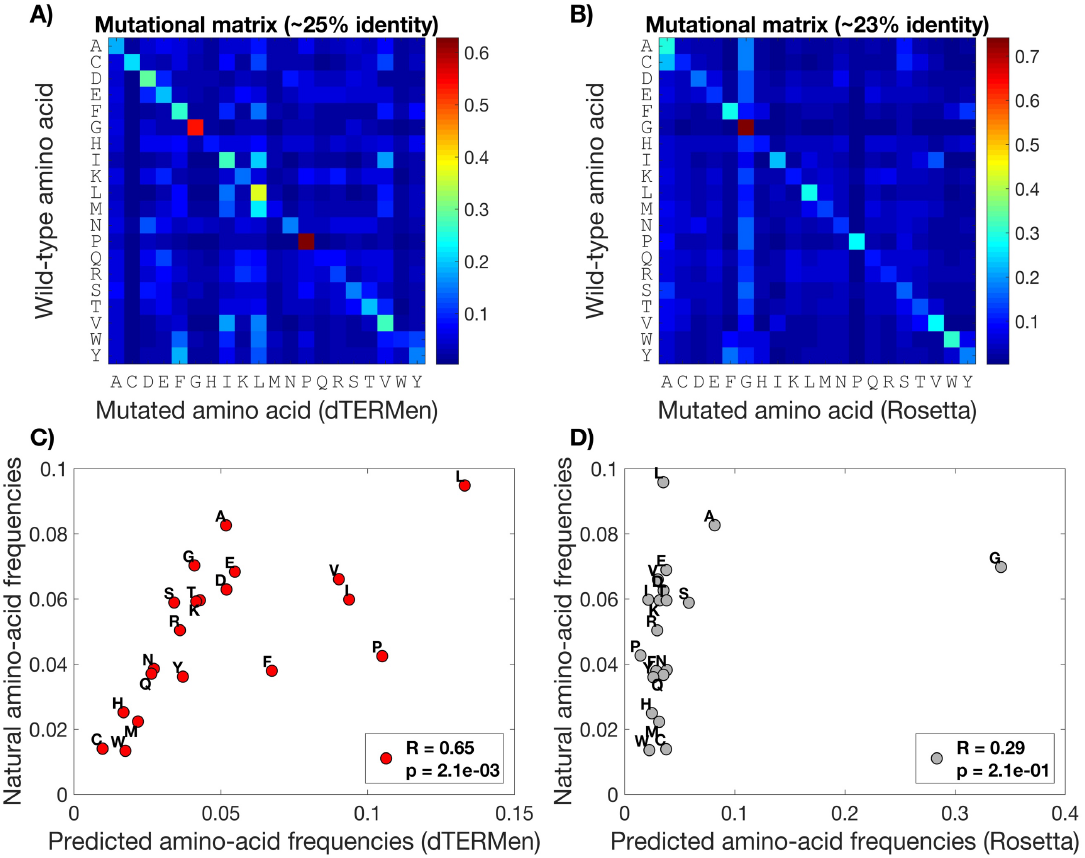
Pattern of amino-acid substitutions predicted by dTERMen is consistent with native amino-acid utilization. Shown in **A)** and **B)** are the mutational matrices predicted by dTERMen and Rosetta Design, respectively. Each entry in the matrix is the conditional probability ***p*(*X*|*Y*)**, as described in the main text, where ***X*** and ***Y*** are the amino acids indicated on the X and Y axes, respectively. Color indicates value in accordance with the show color bar. In **C)** and **D)** the stationary amino-acid distributions implied by the matrices in A) and B), respectively, are plotted against the native amino-acid distribution found in the PDB.

For comparison, we performed the same analysis using Rosetta Design (design procedure shown in Materials and Methods), with the corresponding mutational matrix and emergent stationary amino-acid distribution shown in Figs. 3B,D. Notably, the diagonal elements of *M* are high for both dTERMen and Rosetta, with an aggregate (frequency-weighted) rate of recovering the native amino acid of ∼25% and ∼23% for the two methods, respectively (similar to the values in the native sequence recovery test). However, the stationary distribution implied by the Rosetta matrix indicates clear biases; for example, Gly is expected to be utilized much more frequently than it is in nature. The overall correlation with natural amino-acid utilization is lower than for dTERMen and is not statistically significant.

While both dTERMen and Rosetta choose similarly native-like sequences given native backbones, the underlying model in dTERMen appears more consistent with the natural pattern of amino-acid utilization. This is not entirely unexpected, given that dTERMen designs based on native instances to TERM matches. But the result nevertheless validates the specific statistical approach we use here to extract effective pseudo-energy contributions from structural data.

### 2.5 dTERMen statistical energy indicates design quality

In a recent *tour-de-force* study, Baker and co-workers designed *de novo* and experimentally characterized ∼16,000 sequences for four distinct topologies (see Fig. 4A-D, top) (30). Each design, along with an approximately equal number of negative-control sequences, was tested, in high throughput, for the ability to form folded, stable, protease-resistant structures. These data represent an unprecedented opportunity for testing novel design methods, and we apply them to test dTERMen here. *De-novo* design is a challenging task. So, while each of the ∼16,000 designs represented a sequence predicted to be well compatible with the desired target backbone by the Rosetta Design (2), most designs failed to fold (30). We sought to test whether dTERMen would better distinguish between successful and failed designs. To this end, we ran dTERMen on each of the ∼16,000 backbone structures deposited by Baker and co-workers (one for each of their designs) (30), enabling us to evaluate any natural amino-acid sequence on any of the target models. Next, the dTERMen energy score was computed for each designed sequence on its respective backbone, divided by sequence length to facilitate comparison across different topologies. Fig. 4A-D shows, for each of the four topologies, the correlation between the resulting score and the experimental “stability score”—a protease resistance-based metric the authors developed to estimate design stability in high throughput, having shown it to correlate closely with thermodynamic stability (30). In each case, the correlation is highly statistically significant (p-values in legends; Fig. 4A-D). In contrast, when we considered Rosetta scores computed for each sequence (also published by Baker and colleagues), the correlation is notably weaker in all cases (see Fig. 4E-H), and is either statistically insignificant (p-value of 0.1 in Fig. 4G) or of the wrong sign (positive correlation instead of the expected negative in Figs. 4F,H) in three out of four cases. This result suggests that TERM-based scoring synthesizes structure-sequence relationships in a way that is not easily captured with the existing state-of-the-art in CPD. Further, the ∼16,000 designed sequences were optimized with respect to Rosetta and not TERM-based scoring. In fact, dTERMen best-scoring sequences always differed considerably from Rosetta-based designs (i.e., on average, only ∼16% of positions were identical between Rosetta- and dTERMen-chosen sequences). The fact that dTERMen scores quantify design quality even for sequences that are far from the optimality region of its own predicted sequence landscape validates the generality of the method and the sequence-structure relationships it quantifies. Fig. 5 further shows that the dTERMen score correlates closely with thermodynamic stability, using the same 120 sequence variants of four native domains that Rocklin *et al*. used to establish the quantitative nature of their experimental stability score (30). All of these results together suggest that optimization of dTERMen scores should amount to a robust, general-purpose protein design strategy.

**Fig. 4.**
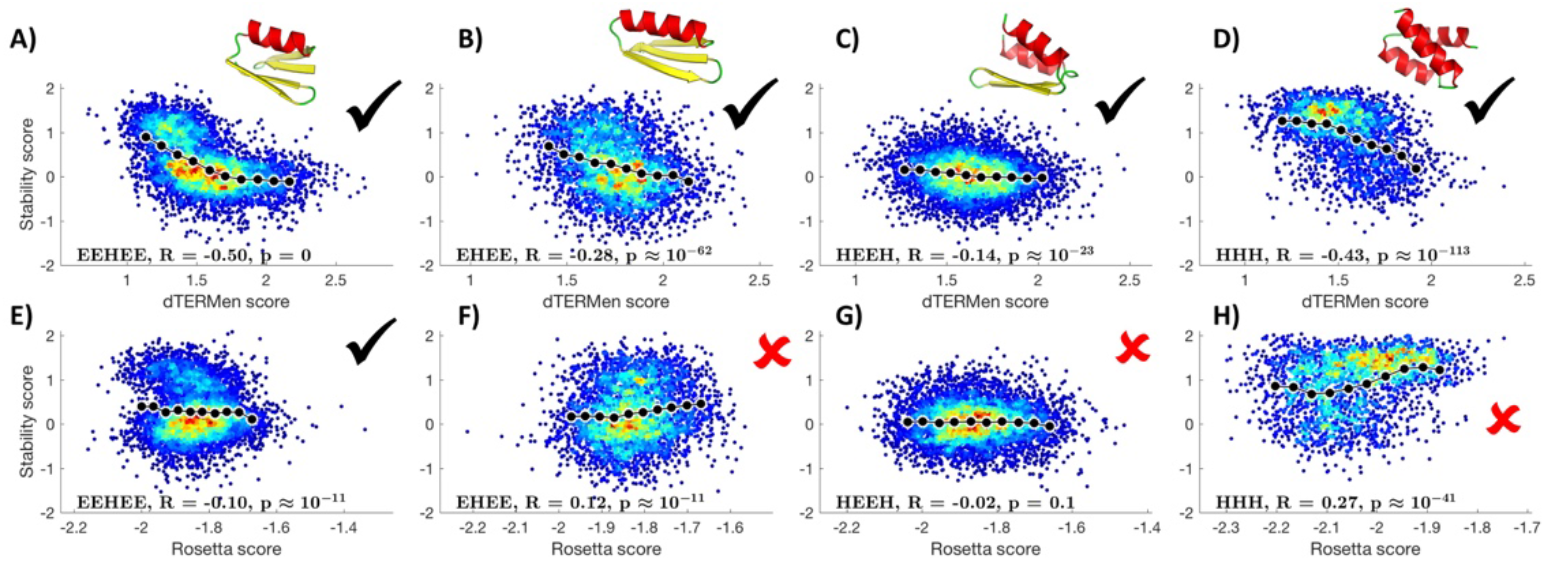
dTERMen statistical energy indicates design quality. Each pair of vertically aligned panels corresponds to one of the four different topologies targeted by Baker and co-workers in their design study (30) (backbone shown above top panel in each pair). Panels **A)**-**D)** show the correlation between the length-normalized dTERMen score of each design (on its respective backbone) on the X-axis and the experimentally-derived stability score for the design on the Y-axis. Point color in the scatter plot indicates data density (decreasing order red-to-blue). The mean curve is shown with a black line with circles, obtained by averaging the stability score in ten progressive windows of the dTERMen score. Panels **E)**-**H)** show the same plots as in **A)**-**D)**, respectively, but with Rosetta score on the X-axis. In each case, the correlation exhibited by dTERMen significantly exceeds that for Rosetta. In fact, in three out of the four cases for Rosetta, the correlation is either of the wrong sign or is statistically insignificant (panels indicated with red X’s), while the correlation is always of the right sign and statistically highly significant for dTERMen (as indicated by black checkmarks).

**Fig. 5.**
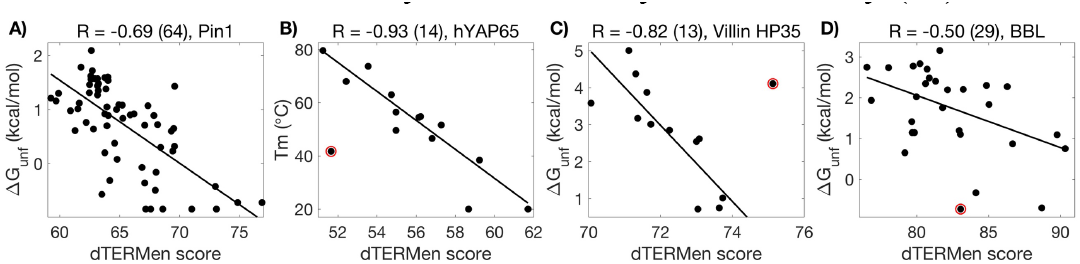
dTERMen scores correlate with thermodynamic stability. Panels **A)**-**D)** correspond to variants of human Pin1 WW domain (modeled via PDB entry 2ZQT), human Yes-associated protein 65 WW domain (modeled via PDB entry 4REX), villin headpiece helical subdomain (residues 42–76; modeled using PDB entry 1VII), and peripheral subunit-binding domain family member BBL (modeled with PDB entry 2WXC), respectively. Each data point corresponds to a single sequence variant, with its thermodynamic stability plotted against its dTERMen score. Thermodynamic stability is represented by the free energy of unfolding in **A), C)**, and **D)**, and apparent melting temperature in **B)**. Best-fit lines are produced using robust linear regression with bisquare weighting function. The Pearson correlation is show in the title for each panel. Outlier points, identified using the Tukey fences approach, are labeled with a red outline and not included in calculating correlation coefficients.

### 2.6 A case study in *de-novo* design by dTERMen

Since dTERMen designs sequences based on information from available native protein structures, would the method still apply if the design target is a *de-novo* generated backbone and not a native one? To interrogate this issue, we considered one of the *de-novo* generated backbones for which Rocklin and co-workers report a successfully designed sequence in their recent large-scale design study (30); see Fig. 6A. Running dTERMen on this specific backbone, letting it choose any natural amino acid at any of the positions (for a total sequence space of ∼10^52^), identifies the solution shown in Fig. 6B as optimal. The modeled structure of the designed sequence looks biophysically reasonable upon close inspection (see Fig. 6B). Further, submitting the designed sequence to HHpred, a powerful structure-prediction method that relies on the ability to identify remote “homologies” between the modeled sequence and a protein of known structure (60, 61), reveals PDB entry 5UP5 as the closest match (with a probability of over 97% and alignment coverage of 90%)—the very experimental structure of the corresponding sequence designed by Rocklin *et al*. (30) (see Fig. 6C). Importantly, 5UP5 was not itself used in the database of proteins from which dTERMen sought TERM-based sequence statistics (and, because it itself is a *de-novo* design, no homologues of it were in the database either). Incidentally, the second match revealed by HHpred, PDB entry 1UTA, is a native structure with a fold highly reminiscent of the target (see Fig. 6D). This is strong evidence suggesting that the dTERMen-designed sequence has the necessary features to be especially favoring of the target structure.

**Fig. 6.**
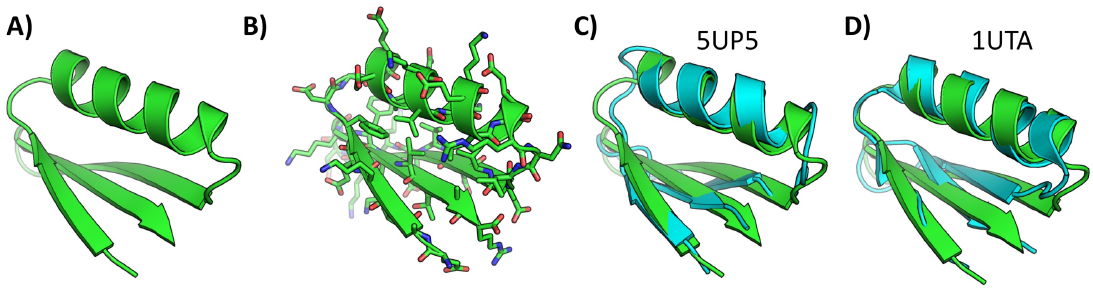
dTERMen can be applied to design structures generated *de-novo*. The backbone shown in **A)** is one of the *de novo*-designed structures targeted by Rocklin *et al*. (30). **B)** Structural model of the sequence designed by dTERMen for this backbone (sequence shown on the bottom). All 40 positions were allowed to take on any natural amino acid. **C)** Superposition between the target backbone (green) and the experimentally-determined structure of the corresponding design by Baker and coworkers (cyan) (30). This structure (PDB code 5UP5) is the top hit for the dTERMen-designed sequence produced by the structure-prediction method HHPred (61). The second hit is the PDB entry 1UTA, whose relevant portion (cyan) is shown superimposed onto the target backbone (green) in **D)**. Sequence of the dTERMen design: eatkefdgpeeaekvkkeleernlevevekkdgkykvtar.

### 2.7 Redesign of mCherry surface

Protein surfaces—i.e., the set of residues exposed to solvent—are important in determining a multitude of biophysical properties, including solubility, immunogenicity, self-association, propensity for aggregation, stability, and fold specificity. It is, therefore, sometimes useful to redesign just the surface of a given protein, so as to modulate one or more of these properties, while preserving its overall structure and function. As an example, let us consider the task of redesigning the surface (resurfacing) of a Red Fluorescent Protein (RFP). RFPs are proteins that naturally fluoresce, with the emission spectrum centered around the red portion of the visible spectrum (∼600 nm). Like other fluorescent proteins (FPs), RPFs are of high utility as biological imaging tags and in optical experiments (62). It may therefore be useful to modulate the surface residues of an RFP depending on the environment (or cell type) in which it has to function.

The crystal structure of RFP mCherry (PDB code 2H5Q (63)) was used as the design template. A total of 64 positions in the structure were manually chosen as being on the surface (corresponding approximately to positions with values of our *freedom* metric above 0.42; see Materials and Methods); these are shown as spheres in Fig. 7A. Following this, dTERMen was used to compute a statistical energy table corresponding to all of the surface positions varying among the twenty natural amino acids, with the remaining positions fixed to their identities in the PDB entry 2H5Q. The resulting energy table, therefore, described a sequence space of 20^64^ ≈ 2·10^83^ sequences. ILP was used to optimize over this space, finding the single sequence with the lowest total statistical energy score. The resulting sequence, compared to the starting sequence of mCherry, is shown in Table IV. The *in-vacuo* surface electrostatic potential of the original mCherry structure and the resulting design model structure are compared in Fig. 7BC. Clearly, the designed sequence represents a significant perturbation to surface electrostatics and overall shape. In fact, a total of 48 out of 64 variable positions are changed in the design.

**Fig. 7.**
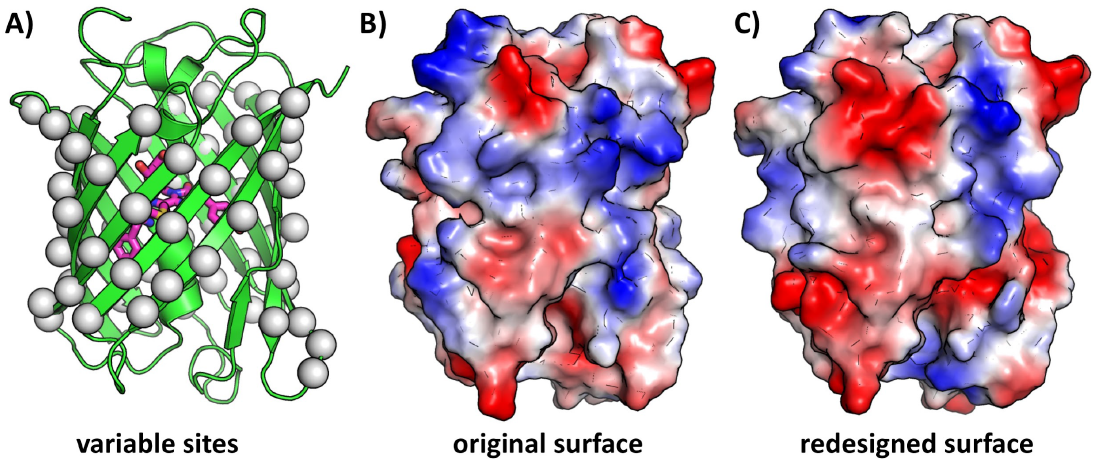
Total surface redesign of mCherry. Show as gray spheres in **A)** are the 64 surface positions that were allowed to vary in design. **B)** and **C)** show the surface of the original mCherry and the redesigned variant, respectively, with the vacuum electrostatic potential designated with false color.

**Table IV.**
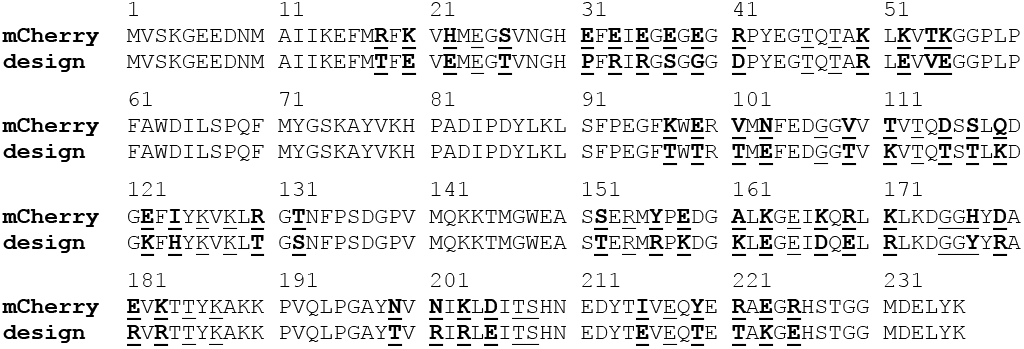
dTERMen design differs significantly from the original wild-type mCherry. Positions marked as variable in design are underlined, and those mutated in the designed sequence additionally marked in bold.

To validate the design, the sequence was cloned into *E. coli*, followed by expression and purification using standard molecular biological and biophysical techniques. Size Exclusion Chromatography (SEC) showed the protein to be monomeric in solution, just as the native mCherry (see Fig. 9A,B), and the far-UV Circular Dichroism (CD) spectrum was consistent with a native-like secondary-structure distribution (see Fig. S1). Despite harboring 48 mutations and despite the fact that preservation of optical properties was not a design constraint (only preservation of structure was), the design still exhibited the chromophore features characteristic of the original protein (see Fig. 9C). Further, the designed protein was still fluorescent, with an emission spectrum of the same shape (but lower intensity) as that of mCherry (see Fig. 9D). Finally, chemical denaturation by guanidinium hydrochloride (GuHCl) revealed that the protein’s structure protects its chromophore approximately as well as the original mCherry—a hyper-stable, highly engineered protein in its own right (Fig. 8). Thus, by all measures, the designed protein, which differs from the original mCherry in 48 positions, still preserves the starting structure and even function. The ability to generate such diversity can be easily exploited to quickly engineer variants of RFP or other proteins that possess a range of desired properties.

**Fig. 8.**
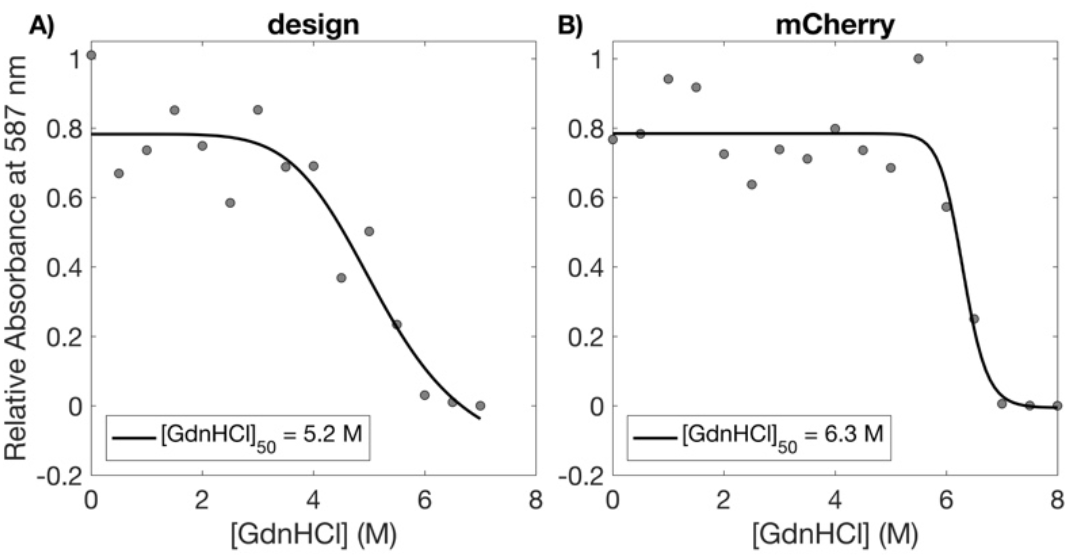
dTERMen-designed variant of mCherry preserves stability. Shown in **A)** and **B)** are chemical denaturation curves, as a function of GdnHCl concentration, of the dTERMen redesign and wild-type mCherry, respectively. Degree of foldedness was monitored via chromophore absorbance at 587 nm. As the chromophore rapidly hydrolyzes upon exposure to water, this constitutes a sensitive metric of structural integrity. Data are fit to the Hill equation, with the apparent concentration of half denaturation noted in the legend.

**Fig. 9.**
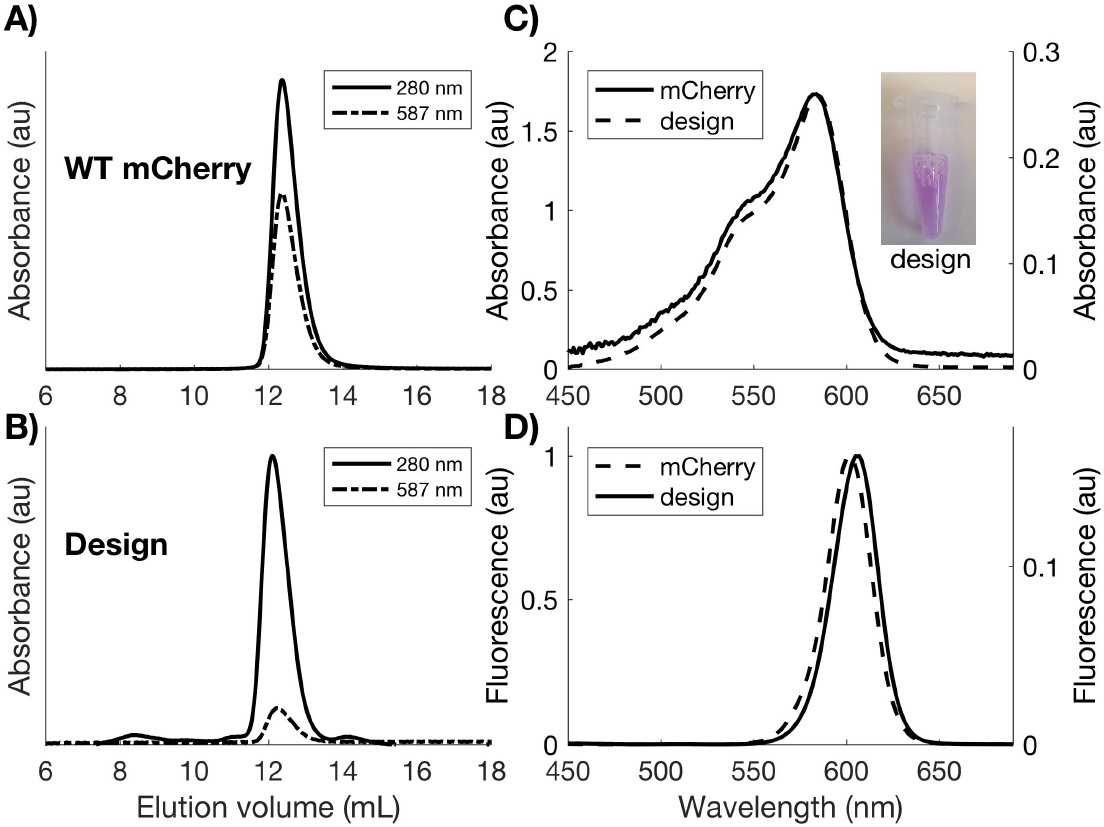
Solution properties of designed mCherry. Shown in **A)** and **B)** are the size-exclusion chromatograms of wild-type mCherry the redesigned variant, respectively, run under identical conditions. The design elutes at a nearly identical volume as the wild type (difference in the ratios between absorbances at 280 nm and 587 nm reflect the lower brightness of the design). **C)** and **D)** further demonstrate that redesigned mCherry preserves photo properties of the wild-type fluorophore. In **C)** absorbance spectra of wild-type and redesigned mCherry are compared (absorbance values shown on the left and right Y-axes, respectively), while **D)** compares fluorescence spectra of the two (left and right Y-axes, respectively). Spectra in both **C)** and **D)** were taken at equivalent concentrations for the two proteins, with Y-axes units reflective relative intensities.

## 3 Discussion

That protein structure can be designed computationally was established a long time ago (1) and has been demonstrated numerous times since (23, 26). It is also true that reliance on prior structural data has been broadly explored, both in terms of various statistics-based methods (46, 49, 52, 54) as well as in the creation of chimeras that fused domains or larger fragments from existing structures (64–67). What is new and exciting about our results here is the marriage between the generality of our approach (i.e., its ability to design sequences for arbitrarily-defined structures) and its reliance on motif-based structural data. This combination is made possible by the fact that protein structure is not “analog” but “digital” in its nature (42, 44, 45)—local-in-space structural motifs, TERMs, tend to be broadly reused across unrelated proteins. These motifs are small enough to be well sampled in the PDB, but large enough to contain non-trivial sequence determinants of structure. One can thus consider an entirely novel structural template as a design target, while still relying purely on existing structures to select sequences well-suitable for folding into it.

Our sequence recovery results, which compare performance on NMR vs. X-ray backbones (see Table I), alternative structures of the same protein (see Table II), and dissect performance by position burial (see Table SI), suggest that, relative to dTERMen, Rosetta derives a lot of insight from geometric fit. This is a consideration that arose as important early in the history of protein design, as researchers observed that X-ray structures generally exhibited jigsaw puzzle-like packed cores (68). But such ideal packing is only feasible in the context of a ground state-like structure. When it comes to room-temperature ensembles, the requirement for a crystalline packed core may not be appropriate. Backbone flexibility techniques have been proposed to address the issue that while rotamer-based methods are effectively modeling the ground state, a pre-specified template may not represent such a state for any designed sequence (9, 69). In dTERMen this issue is address implicitly, to an extent, by the fact that sequence statistics are gathered from ensembles of close TERM matches. Our extensive tests here (from native sequence recovery, Tables I–II, to the prediction of design success and thermodynamic impacts of mutations, Figs. 4–5, to a *de-novo* design example, Fig. 6, and the redesign of mCherry, Figs. 7–9) suggest that this does appear to be a valid way of dealing with the issue. But more work is needed to identify the best means of representing the ensemble nature of structure while data mining in the context of TERMs.

In a traditional atomistic approach to design, specific important aspects of the physics underlying protein structure are recognized, parameterized, and included as part of the scoring function. Then, sequences are chosen based on this quantitative (albeit highly approximate) model. In dTERMen, the fundamental “reasons” behind sequence choice are not described beyond observed biases in sequence distributions among database sub-structures. This can be seen as a disadvantage. The corresponding advantage, however, is that complex effects can be included without the need for understanding of their origin (or even being aware of them). A good example of this is disulfide bonds. The physics of these covalent bonds between side-chains of cystine residues is not trivial to model, and to strike the right balance between when to include such bonds and when not to in designing proteins is also not easy. However, as shown in Table I, dTERMen frequently places disulfide bonds at locations where they appear natively. Importantly, this is not because dTERMen chooses cystines too often—in fact, Cys occurred at a frequency of ∼1% in dTERMen-designed sequences compared to the rate of ∼2.5% within corresponding native sequences. And, in general, amino-acid utilization implied by the dTERMen model is in good agreement with the native distribution of amino acids (see Fig. 3).

When inspecting design models manually, we frequently see other examples of dTERMen automatically recognizing and utilizing well-known sequence-structural patterns, such as helix-capping motifs (70), salt-bridge patterns within different secondary-structure combinations (71), b-turn preferences (39, 72), and even p-cation interactions (73). But these are just some of the patterns that we recognize, based on our experience with protein structure. It is interesting to consider what other important sequence-structure patterns—those *not* already well known—may be automatically included in dTERMen designs.

In summary, the evidence presented here strongly points to the fact that design of protein structure based on sequence patterns mined from the PDB is feasible and practical. Based on extensive benchmarking, our general-purpose design framework dTERMen performs on par with or better than the state of the art in CPD. What is most exciting about this finding is that the “top-down” TERM-based insights that dTERMen relies upon are quite distinct from to the “bottom-up” MM-based models that are typically used in CPD. We can thus reasonably expect that the two methodological classes will have orthogonal strengths and weaknesses. There should be ample opportunity to improve the overall robustness of CPD as a whole by combining TERM- and MM-based insights and by further optimizing the specifics of TERM-based structure mining.

## 4 Code availability

dTERMen is implemented as a python-based program that makes extensive use of our structure search engine MASTER (43) to identify TERM matches and extract their sequence statistics. The source code for dTERMen can be found at https://grigoryanlab.org/dtermen.

## 5 Acknowledgements

This work was funded by NSF award DMR1534246 (GG) and NIH award P20-GM113132 (GG).

## 6 Materials and Methods

### 6.1 Pseudo-energy computation and optimization

#### 6.1.1 Pre-computed contributions

The pseudo-energy contribution describing the propensity of different amino acids for backbone *φ* and *ψ* dihedral angles, perhaps the best understood structural preferences in proteins, is the first one in our hierarchy. This is a traditional statistical potential, built by splitting the *φ*/*ψ* phase-space into bins (e.g., bins of 10° × 10°) and assigning each residue in a database of native proteins (the “native database”) into a corresponding bin based on its *φ*- and *ψ*-angle values. Then, the pseudo-potential for amino acid *a* associated with backbone dihedrals bin 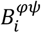 is computed as: 
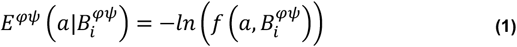
where 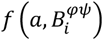 is the frequency with which amino acid *a* is found in this bin within native proteins: 
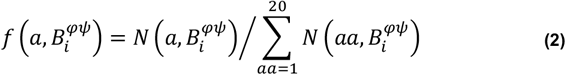
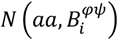 being the number of times amino acid *aa* is found in bin 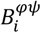. The next contribution in our hierarchy is associated with the preference of amino acids for different backbone *ω* dihedral angles. Because this angle is defined around the peptide bond, which has partial double-bond character, *ω* angles are typically planar, with values close to 180° most common (trans peptide bonds), but values around 0° also occurring (cis peptide bonds), generally (though not exclusively) with Pro or Gly amino acids. Other values also occur, though infrequently. For this reason, we choose a nonuniform binning of *ω* angles, where bin widths are at least 1°, but as large as needed to have a sufficient number of database residues in each bin. Then, the pseudo-energy of amino acid *a* associated with *ω*-angle bin 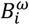 is calculated as: 
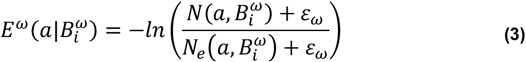
where 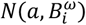 is the number of times amino acid *a* is found in bin 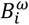, and 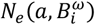 is the number of times *a* is expected to be found in the bin, based on the pseudo-energy contributions already known—namely, the *φ*/*ψ* energy. The added value *ε*_*ω*_ acts as a pseudo-count, preventing excessive statistical noise from poorly populated bins (we generally use *ε*_*ω*_ = 1). The expectation where expectation 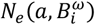 is calculated as: 
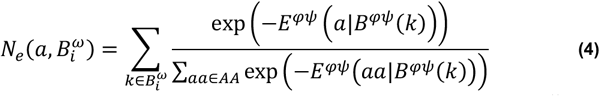
where the outer sum is over all native residues falling into *ω* bin 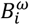, the inner sum is over all natural amino acids, denoted by set *AA*, and *B^φψ^*(*k*) is the *φ/ψ* bin bin into which residue *k* falls. The inner fraction represents the expected probability of observing *a* (over all possible amino acids) in the *φ*/*ψ* environment of each residue in the bin. The correction by expectation in Eq. 3 assures that *E^ω^* acts only as a corrector over *E^φ/ψ^*, explaining only what is not already explained in the data. We adhere to this principle in building the rest of the pseudo-energy hierarchy.

The next contribution we consider is that from the general environment (burial state) of a residue. To capture this as a single-body (self) contribution, one needs a metric describing the environment of a residue in a protein structure based solely on the backbone configuration (i.e., independent of any specific sequence associated with the structure). This is not trivial, because if we consider the burial state of a residue *per se* (e.g., via solvent-accessible surface area), it will be highly sensitive to the amino acid occupying the residue as well as the surrounding sequence. To meet the challenge of a sequence-independent environmental descriptor, we propose the metric we call residue *freedom*, which considers all possible rotamers of all natural amino acids at a given position and its surroundings to determine the extent to which the volume around the residue would tend to be unoccupied and available to its rotamers (see “Methodological Details” for exact definition). Freedom has worked well for us in practice, though any sequence-independent environmental descriptors can be used in its place. Whatever the descriptor, we will call it *environment* and will denote it as *e*. To build an environmental pseudo-potential, we compute *e* for all residues in the database and bin them according to this value (freedom values range from 0 to 1, and we uniformly bin them into ∼70 bins). Then, the pseudo-energy of each amino acid *a* associated with environment bin 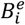 is pre-computed as: 
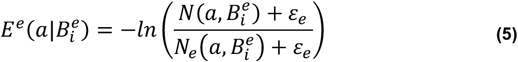
where 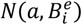 is the number of times amino acid *a* is found in bin 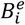, and 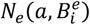 is the number of times *a* is expected to be found in the bin, based on the pseudo-energy contributions already known—*φ*/*ψ* and *ω* energies. The value *ε^e^* acts as a pseudo-count, preventing excessive statistical noise from poorly populated bins (we generally use *ε^e^* = 1). The expectation 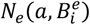 is calculated as: 
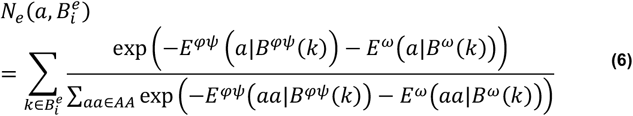
where the outer sum is over all native residues assigned to the environment bin 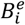, and *B^ω^*(*k*) is the *ω* bin into which residue *k* maps. Here again, the environment potential acts as a corrector over what is already explained by *φ*/*ψ* and *ω* pseudo-energies.

#### 6.1.2 TERM-based contributions

The above potentials are all pre-computed ahead of time, based on the native database. The following several pseudo-energy correctors in the hierarchy are EPs that need to be computed on-the-fly, while performing design calculations, by analyzing TERMs and their matches. The first one is the own-backbone correction, which captures how the local contiguous stretch of backbone around position *p* modulates its amino-acid preferences, beyond what is already captured by *φ*/*ψ*/*ω* preferences. To this end, we define a singleton TERM around *p*, lets call it *T_p_*, finding all of its non-redundant close structural matches in the database. Lets refer to this set of matches as *M_p_*. Then, the own-backbone energy of amino acid *a* in position *p* is calculated as: 
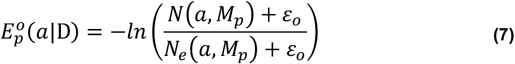
where *N*(*a, M_p_*) is the number of times we observe *a* in the position corresponding to *p* within TERM matches *M_p_* and *N_e_* (*a, M_p_*) is the number of times we expect to see *a* in this position, based the pseudo-energy model available so far (i.e., *φ*/*ψ, ω*, and environment energies). The value *ε_0_* is a pseudo-count, and we generally use the value of 1. By analogy to previous instances, the expectation in the denominator is computed as: 
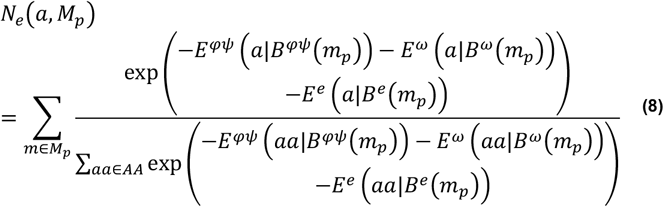
where the outer sum is over matches in *M_p_, m_p_* is the residue in match *m* that aligns with position *p* in *T_p_*, and *B^e^*(*m_p_*) is the environment bin to which *m_p_* belongs, based on its surroundings in the structure from which match *m* originates.

The next contribution in the hierarchy is the near-backbone correction. It captures any further modulation of amino-acid preferences at *p* brought about by the presence of backbone segments in close spatial but not sequence proximity to *p*. This contribution is deduced from matches to near-backbone TERMs, as described in “TERM decomposition” above. Ideally, one would have a single such TERM that covers all proximal backbone segments. But due to a limited database size, such a TERM may which exhibit an insufficient number of close structural matches (and thus poor sequence statistics), so multiple near-backbone TERMs may need to be used to cover sub-sets of proximal backbone segments (see “TERM decomposition”). For each near-backbone TERM considered around position *p*, 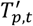 (subscript *t* indicates that multiple such TERMs are possible), we compute a set of amino-acid correction pseudo-energies by considering the non-redundant close structural matches to the TERM. Referring to this set of matches as 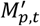, we compute the near-backbone correction associated with amino acid *a* at TERM *T_p,t_*: 
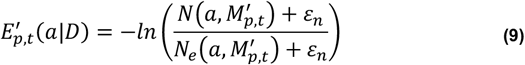
where 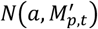 is the number of times we observe *a* in the position corresponding to *p* within matches 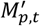 and 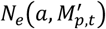 is the number of times we expect to see *a* in this position, based the pseudo-energy model available so far (i.e., *φ*/*ψ, ω*, environment, and own-backbone energies). The value *ε_n_* is a pseudo-count (see section “Pseudo-counts” for details). The expectation in the denominator is computed as: 
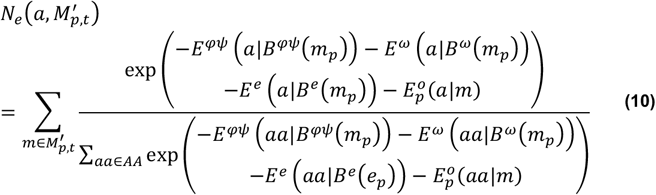
where the outer sum is over matches in 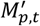 and 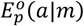 represents the own-backbone correction for amino acid *a* in residue *m_p_*, based on the structure from which match *m* originates. Note that to correctly compute 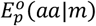, one must evaluate Eq. 7 for each match in 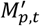. This means searching for singleton TERM matches defined around *m_p_* for each match in 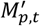. This is because one cannot simply assume that the own-backbone correction for the original residue *p* will be the same as that for the matching residues *m_p_* in near-backbone TERM matches. In practice, this computation of 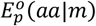 takes a considerable additional time and, often, leads to relatively small corrections. For this reason, we typically disregard 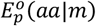 when computing the expectation *N_e_* 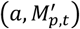. This can lead to some over-counting of own-backbone effects, as they would be partially reflected in both own-backbone and near-backbone corrections, but in our experience this over-counting is minor. Further, own-backbone effects, being low order, are statistically more robust than higher-order corrections, so some over-counting of the former in the final model may be tolerable.

##### 6.1.2.1 Pair contributions

Next in our hierarchy are amino-acid pair pseudo-energies. These are defined for each pair of coupled positions in *D*, (*p, q*). For each such pair, as described in “TERM decomposition”, we generate a pair TERM, lets call it 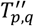, with a set of non-redundant structural matches, 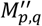. From these, the pseudo-energy pair potential between amino acids *a* and *b* in positions *p* and *q*, respectively, is computed as: 
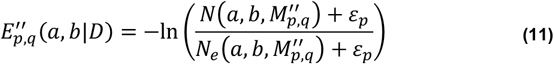
where *N*(*a, b*, 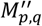) is the number of times we observe *a* and *b* in the positions corresponding to *p* and *q*, respectively, within matches 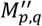 and *N_e_*(*a, b* 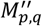) is the number of times we expect to see (*a, b*) pairs in these positions, based the pseudo-energy model available so far. The value *ε_p_* is a pseudo-count (see section “Pseudo-counts” for details). *N_e_*(*a, b*, 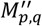) is computed as: 
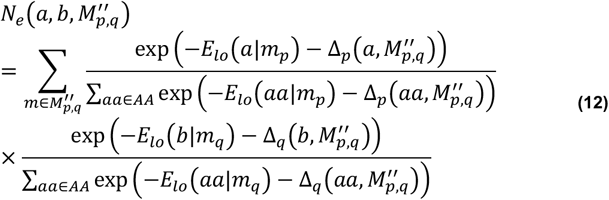
where for brevity we denote with *E_ιo_*(*a*|*m_p_*) the total pseudo-energy, from all lower-order contributions considered thus far, associated with amino acid *a* in the position aligned with *p* of match *m*: 
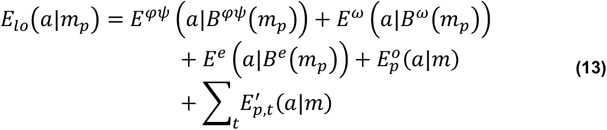
and Δ_*p*_ (*a*, 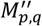) is an adjustment energy that can be included to preserve the marginal amino-acid distributions at individual coupled positions of the pair TERM. Introducing this marginal constraint is optional, but advantageous in our experience. This is because at the point in the hierarchy when we compute pair contributions, we expect all (or nearly all) of the significant effects on the marginal amino-acid distribution at individual positions to have been captured by lower-order energies, with any remaining unaccounted positional preferences likely to be due to statistical noise. Nevertheless, with sufficient amounts of structural data, this constraint should become unnecessary and the adjustment energy can be omitted. Note that even when included, the marginal constraint still allows the pseudo-energy model to dictate how pairs of amino acids across the two positions are expected to distribute. The adjustment energy is computed as: 
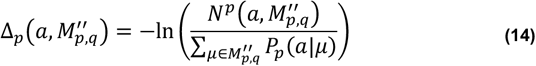
where *P_p_*(*a*|*μ*) is the unadjusted marginal expected probability of observing amino acid *a* in the position aligned with *p* within match *μ* and *N^p^*(*a*, 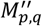) is the number of times *a* is observed in this position across all matches in 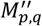. The ratio under the logarithm is between the expected and observed number of times *a* is seen in position *p*. The un-adjusted marginal expectations at each position are computed as: 
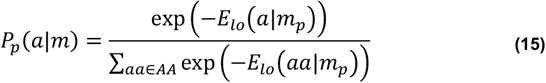

Note again that to evaluate in Eq. 13 fully, the computation of 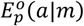 and 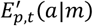, for each match *m* and each near-backbone TERM that it may give rise to within its parent structure, is computationally quite costly. Thus, we generally omit these pseudo-energy components when computing *E_ιo_*(*a*|*m_p_*).

##### 6.1.2.2 Higher-order contributions

The framework generalizes to allow the computation of higher-order interaction EPs, although sequence statistics of structural matches quickly become limiting for these contributions and, in practice, we typically do not go beyond pair components. Because higher-order contributions are corrections on pair- and lower-order ones, it is not unreasonable to assume that they vanish. Still, for completeness, we present an example of how one might compute triplet pseudo-energy corrections. These could be defined for any three positions (*p, q, r*) in *D* that are a connected sub-graph of *G* (i.e., they can each influence each other, directly or indirectly; see “TERM decomposition” for details). The corresponding triplet TERM, 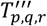, is constructed as detailed above and its non-redundant close structural matches obtained, 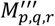. Following this, the pseudo-energy triplet potential between amino acids *a, b*, and *c* in positions *p, q*, and *r*, respectively, is computed as: 
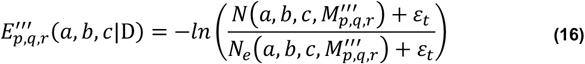
where *N*(*a, b, c*, 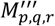) is the number of times we observe amino acid combination (*a, b, c*) in positions (*p, q, r*), respectively, within TERM matches 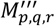 and *N_e_*(*a, b, c*, 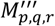) is the number of times we expect to see (*a, b, c*) in these positions, based the pseudo-energy model available so far. The value *ε_t_* is a pseudo-count (see section “Pseudo-counts” for details). *N_e_*(*a, b, c*, 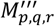) is computed as: 
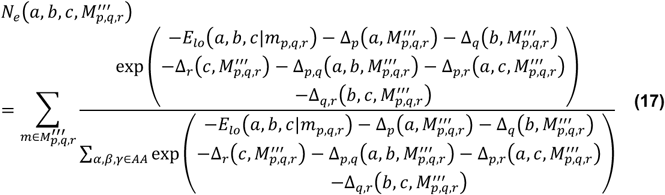
where *E_ιo_*(*a, b, c*|*m_p,q,r_*) denotes the total pseudo-energy, from all lower-order contributions considered thus far, associated with amino acids *a, b*, and *c* in the positions *p, q*, and *r*, respectively, of match *m*: 
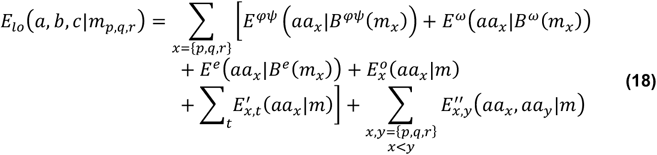
and Δ*p,q*(*a, b*, 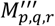) is an optional pair adjustment energy, analogous to the marginal adjustment introduced when computing pair interactions, meant to constrain the pairwise amino-acid distributions at pairs of positions in 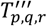 to the observed distributions. The argument here is similar—pair preferences should have been captured previously in pair pseudo-energies, such that any additional pair preferences in 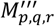 are likely the result of statistical noise. And again, with enough structural data, this adjustment should become unnecessary. Marginal adjustments are computed as before (see Eq. 14), and pair adjustments as follows: 
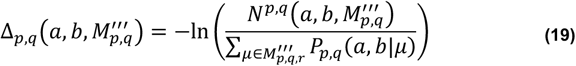
where *P_p,q_*(*a, b*|*μ*) is the expected probability of observing amino acids *a* and *b* in positions *p* and *q* in match *m*, respectively (without the pair adjustment), and *N^p,q^*(*a, b*, 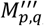) is the number of times *a* and *b* are observed in these positions across all matches in 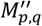. The expected probability without the pair adjustment is calculated as: 
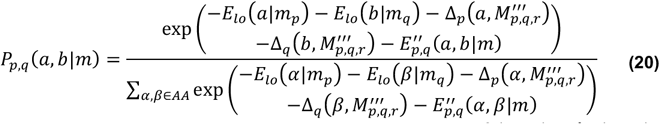

Note that to evaluate Eq. 17 fully, the computation of 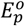(*a*|*m*), 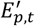(*a*|*m*), and 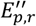(*aa_x_, aa_y_*|*m*) for each match *m* and each near-backbone TERM that it may give rise to within its parent structure, is computationally quite costly. Thus, we generally omit these pseudo-energy components when computing *E_ιo_*(*a, b, c*|*m_p,q,r_*).

#### 6.1.3 Coupled residues

Presently, we use the metric of *contact degree*, introduced in our previous studies (42), to designate coupled positions in a given structure. The contact degree between positions *i* and *j* is computed as: 
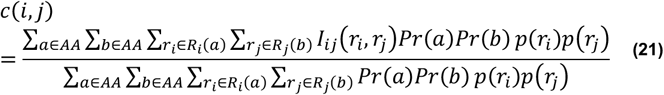
where *R_i_*(*a*) is a set of side-chain rotamers of amino acid *a* at position *i* (after discarding rotamers that clash with the backbone), *I_ij_*(*r_i_, r_j_*) is a binary variable indicating whether the two rotamers *r_i_* and *r_j_* would likely strongly influence each other’s presence (have non-hydrogen atom pairs within 3 Å), *Pr*(*a*) is the frequency of amino acid *a* in the structural database, and *p*(*r_i_*) is the probability of rotamer *r_i_*. Rotamers and their probabilities can be taken from any backbone library, but we have generally used the backbone dependent library developed by Dunbrack and coworkers (10). By construction, the value *c*(*i, j*) varies between 0 and 1, with higher numbers corresponding to position pairs that are more poised to influence each other. Thus, one can use a given contact-degree cutoff to identify which position pairs are to be considered coupled for the purposes of design calculations. At present, we use the cutoff of 0.001.

#### 6.1.4 Environment metric

We currently use the metric of *freedom*, which we previously introduced, to quantify the general degree of burial of a given position, although our framework can be used with any sequence-independent measure. Freedom measures the degree to which side-chain rotamers at a given position are likely to be unobstructed by other side-chains or the backbone, and is calculated as: 
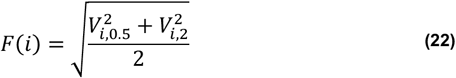

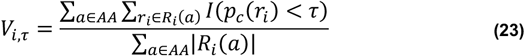

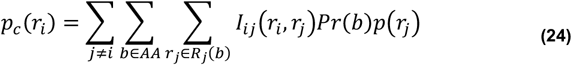
where definitions of *r_i_, R_i_* (*a*), *Pr*(*a*), *p*(*r_i_*), and *I_ij_*(*r_i_, r_j_*) are as in Eq. 21. *p_c_*(*r_i_*) is the “collision probability mass” of rotamer *r_i_*—i.e., how likely it is to clash with rotamers at other positions. Higher *p_c_*(*r_i_*) values indicate that *r_i_* is expected to be generally crowded out by other side-chains in its environment. *V_i,τ_* then quantifies the total weight of rotamers at position *i* that are not overly crowded (i.e., have collision probability masses below *τ*), Finally, we combine *V_i,τ_* with two thresholds *τ*, 0.5 and 2, into the final freedom metric at position *i, F_i_*.

#### 6.1.5 Selection of matches

As described in “TERM decomposition”, every TERM is formed around a set of one or more residues that represent a connected sub-graph of *G* (the graph of residue couplings). Let’s refer to this set of residues as *g* (source residues) and the resulting TERM as *T_g_*. To identify a set of diverse close structural matches to *T_g_*, we consider backbone sub-structures from the native database that align onto the backbone of *T_g_* with low root-mean-square-deviation (RMSD; hydrogen atoms are excluded when calculating RMSD). Our search engine MASTER identifies such sub-structures ordered by increasing RMSD (43). To select final matches, we perform the following procedure for every match *m*, in low-to-high RMSD order, until a certain number *N* of matches are accepted (*N* can be chosen differently based on what pseudo-energy component is being calculated; see below) or until the RMSD exceeds a certain threshold (we employ a size/complexity-dependent cutoff function; see below), whichever comes last. For each residue *r* in *m* aligned to one of the source residues, we extract the local sequence window by considering residues in the range [*r* − 15; *r* + 15] in the structure from which *m* originated. Next, we compare *m* to each of the previously accepted matches by aligning the corresponding sequence windows via Needleman-Wunsch algorithm and the BLOSUM62 matrix. If for any previously accepted match *μ*, any pair of windows produces an alignment with a p-value lower than 10^−6^, then the new match *m* is considered redundant with respect to *μ* and is discarded. Alignment p-values are computed based on alignment scores and indicate the probability that an alignment between sequences of the same length (chosen with database amino-acid frequencies) scores as well or better.

##### 6.1.5.1 Number of matches

A minimal number of TERM matches is needed to derive stable statistics and avoid small-number noise. Depending on the value to be estimated, this number can be chosen differently. At present, we use *N* = 1000 for all pseudo-energy calculations, except for the near-backbone component, where the value of *N* = 200 is used.

##### 6.1.5.2 RMSD cutoff

RMSD is a good relative metric of similarity between TERMs and their potential structural matches. However, an absolute cutoff for defining a match is not trivial to define, because tolerable RMSD values tend to grow with structure size. To address this issue, we previously developed a size- and complexity-dependent RMSD cutoff function for a TERM (42): 
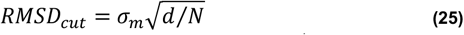

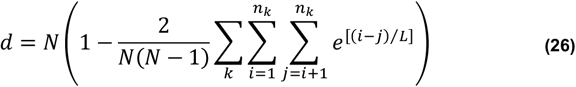
where *d* is the effective number of degrees of freedom for the TERM, *n_k_* is the length of the *k*-th contiguous segment of the motif, *N* is the total length of the motif (i.e., *N* = ∑*_k_ n_k_*), *L* is correlation length—a parameter describing the extent of special correlation between residues in the same polypeptide chain, and *σ_m_* is a plateau parameter. We currently use *L* and *σ_m_* of 20 and 1.0 Å, respectively (and 15 and 1.1 Å for near-backbone searches).

##### 6.1.5.3 Native database

The structural database for use in dTERMen was generated by considering the weekly *BLASTclust* clustering of all PDB chains (as of October 22, 2014) with a sequence identity cutoff of 30% (74). Chains identified as the first element of each resulting cluster were collected and filtered to include only structures resolved by X-ray crystallography to a resolution of 2.6 Å or better; chains with only Cα backbone traces were excluded. This gave the final database of 14,546 protein chains.

##### 6.1.5.4 Pseudo-counts

An appropriate choice of pseudo-count magnitude is key for extracting maximally useful information from sequence statistics. Overly low pseudo-count values tend to produce noisy pseudo-energies, whereas overly high values lead to over-dampened pseudo-energies, such that many important structural effects are not appropriately recognized. Either case leads to poor amino-acid sequences chosen for the target structure. In our formulation, pseudo-counts are added to both observed and expected amino-acid counts, to dampen statistical noise present in both. And we have noticed that best pseudo-count values to use generally depend on the number of expected and observed counts. Thus, we have come up with an empirical formula for the pseudo-count: 
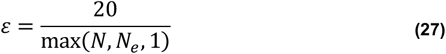
where, *N* and *N_e_* are the observed and expected counts (of whatever quantity is being modeled), respectively. By decreasing with increasing amount of data, this pseudo-count function balances well between capturing important effects accurately, while dampening low-count observations to minimize the effect of noise.

#### 6.1.6 Optimization

Once all EPs are computed and organized into a table of self, pair, and possibly higher-order pseudo-energies, a host of optimization approaches can be used to deduce the optimal amino-acid sequence. We use an Integer Linear Programming (ILP) approach, which conveniently allows us to introduce constraints into the design problem (i.e., sequence symmetry constraints, or constraints on the number of charged/polar residues, or limits on the residues mutated relative to some starting sequence, etc.) However, if ILP fails to converge quickly, any number of alternative optimization methods can be used—e.g., Self-Consistent Mean Field (SCMF) or Simulated Annealing Monte Carlo (MC). Importantly, identification of the absolute global optimal sequence is not strictly required, and any close-to-optimal sequence would suffice.

### 6.2 Native sequence recovery

To arrive at the list of templates used in native sequence recovery tests, the full list of domains in the CATH database (version 4.2.0) was downloaded on Feb 11, 2018 (75). The list was filtered using the following criteria: a) each domain had to be an entire chain of the corresponding PDB entry that was non-deprecated and monomeric (i.e., both biological and asymmetric units containing a single chain), b) domains in the “few secondary structures” CATH class were excluded, c) domains corresponding to membrane-protein PDB entries (i.e., those listed in the OPM database (76)) were excluded, d) only domains ranging from 50 to 150 residues in lengths, consisting entirely of natural amino acids (including MSE, HSC, HSD, HSE, and HSP), and with no missing non-hydrogen backbone atoms were allowed, and e) for X-ray PDB entries, only those with resolution of 2.6 Å or better were allowed. The resulting list was split into 10 bins by domain length (i.e., [50, 60), [60, 70), [70, 80), …, [130, 140), and [140, 150]), with 8 X-ray and 2 NMR structures selected from each bin manually, making sure that structures chosen from the same bin belonged to different CATH topologies. This gave a set of 100 monomeric, single-domain, water-soluble structures with 80 X-ray and 20 NMR entries (set I, see Table SII).

We also considered the 11 pairs of structures, each pair representing one NMR and one X-ray structure of the same protein, curated in our earlier work (i.e., sets X-ray-2 and NMR-2 from MacKenzie *et al*. (42)), here referred to as set II (see Table SIII). Tests comparing design performance on NMR vs. X-ray structures or alternative NMR models used set II structures, while all other sequence-recovery tests used the union of set I and set II, containing 90 X-ray and 31 NMR structures (X-ray entry 1TTZ occurred in both set I and set II). Matching entries 3IBW (X-ray) and 2KO1 (NMR) from set II are homodimers (all others being monomeric), so one of the monomers was kept at its wild-type sequence during design with both dTERMen and Rosetta. Each NMR entry in set II contained 20 models, so each gave rise to 190 model-to-model comparisons, giving a total of 2,090 such comparisons across set II. For NMR-to-X-ray comparisons, each X-ray entry was compared to each of the 20 models of the corresponding NMR entry.

Positions were classified into surface, interface, and core using solvent-accessible surface area (SASA) values computed in the context of the native protein used as the template. Specifically, Stride (downloaded on June 24, 2018) was used to calculate absolute SASA values for each residue (77) and these were divided by “standard” reference SASA values for each amino acid type to obtain relative SASAs. Standard values were taken from GetArea (i.e., for each residue type X, these area the surface area in the tripeptide Gly-X-Gly, averaged over a set of 30 random conformations) (78). Residues were labeled as surface if the relative SASA exceeded 40%, as core if the value was below 20%, and interface for cases between 20% and 40%.

Disulfide bonds in native structures were identified as instances of two cystine residues with SG atoms within 3.0 Å of each other, resulting in a total of 80 and 7 Cys-Cys bonds in all X-ray and NMR structures considered, respectively. Disulfide bond recovery was computed as the fraction of times the designed sequences retained two cystines at position pairs that were natively disulfide bonded.

### 6.3 Design with Rosetta

We used pyRosetta (Linux release r56316.64Bit) in all Rosetta Design tests, as well as to repack dTERMen-designed sequences onto target backbones (e.g., for visualization in Fig. 7). Specifically, we performed fixed-backbone design using the *talaris2013* force-field and default parameters in pyRosetta via “standard_packer_task” and “PackRotamersMover” objects (for building structural models of dTERMen designs, only the single amino acid from the designed sequence was allowed at each position). Specifically, the relevant portion of python code we used is shown in Table SIV.

### 6.4 Structure prediction test

Sequences designed for the 100 structures in set I (see Table SII), as well as their native counterparts, were subjected to structure prediction using standalone I-TASSER (version 5.1, downloaded on June 4, 2018) (79). Specifically, I-TASSER was run in fast mode, with at most 5 hours for each round of simulation, producing at most 10 final models. Information from homologues of the protein used as the design template (i.e., the native sequence) was excluded from I-TASSER prediction. To this end, we used *blastpgp* from the standalone BLAST packages (version 2.2.26) to search the PDB (i.e., the preformatted BLAST database file *pdbaa* downloaded from NCBI on June 6, 2018) for homologues of the native sequence using the E-value cutoff of 1 (74). Chains corresponding to any matches, as well as the design template itself, were then removed from the I-TASSER template library during runs using the `–temp_excl` flag. With these settings, an I-TASSER run took around ∼20 hours wall-clock time, on average.

All the predicted models were further compared to their respective design templates via TM-score and secondary structure recovery. The former was calculated using TM-align (downloaded on June 24, 2018) (53). Stride (downloaded on June 24, 2018) was used to identify the secondary structure for each residue in models and native structures (77).

While I-TASSER was asked to return up to 10 best models for each sequence, fewer models were produced in some cases (due to the inability of the method to identify a sufficient number of structural templates). When comparing models across different sequences categories (e.g., in Fig. 2 and Table III), the same index model was always compared. For example, model 3 for the dTERMen sequence designed on the backbone of protein *X* was compared with model 3 of the Rosetta sequence designed on this backbone. Thus, if (for example) the dTERMen sequence resulted in 10 models and the Rosetta sequence produced 9 models, only the first 9 were compared. In total, there were 481 models that were successfully produced for all three sequence types (dTERMen, Rosetta, and native), and all comparisons were made using only these. This included models for 83 targets (I-TASSER produced no models in 13/100, 13/100, and 15/100 cases for dTERMen, Rosetta, and native sequences, respectively).

### 6.5 Experimental characterization of mCherry design

Both wildtype and design mCherry construct genes were synthesized, sequence-verified, and cloned into plasmids by Gen Script (pUC57 for wildtype mCherry and a modified pET28b for the design variant; in this modified plasmid, the Factor Xa cleavage site was replaced with a Tobacco Etch Virus or TEV protease site). Wildtype mCherry was subcloned into a standard pET28b using Agilent’s QuikChange Lightning site-directed mutagenesis kit through a PCR insertion method relying on distal end homology between insert and template. The cloned construct sequence was confirmed by DNA sequencing (Dartmouth College Molecular Biology Core Facility).

#### 6.5.1 Protein expression and purification

Both proteins were expressed in *E. coli* Rosetta 2 (DE3) cells made competent in house. Expression was carried out through induction for 17 hours at 20°C by addition of 0.2 mM IPTG at an OD_600_ around 0.7–0.9. Cells were subsequently harvested by centrifugation at 3,000 rpm for 25 minutes and the pellets were resuspended in 30 mL FPLC binding buffer (50 mM Tris-HCl, 250 mM sodium chloride, 20 mM imidazole, pH = 8.0). Cells were lysed using a Microfluidizer®. The soluble protein fraction was cleared by centrifugation at 20,000 rpm for 40 minutes. The proteins from the lysed cultures were purified by means of affinity chromatography on a GE Healthcare Akta PureM FPLC system on Ni-NTA conjugated resin (GE Healthcare HisTrap HP 5 mL column) followed by size exclusion chromatography (SEC) on a GE Healthcare Superdex-75 16/600 prep-grade column or a Superdex Increase 10/300 GL column.

#### 6.5.2 Circular dichroism

Folding and stability of wildtype and design mCherry constructs were assessed by circular dichroism (CD) on a Jasco J-815 instrument. All samples contained 10–20 μM protein in 25 mM sodium phosphate, 150 mM sodium chloride, pH = 7.5. CD scans were acquired at 20°C with four accumulations each in the 250–200 nm UV range, at 100 nm/min, and with a 1 nm bandwidth, and a pitch of 0.1 nm.

#### 6.5.3 Fluorescence

Fluorescence spectra were recorded on a synchronous scanning Jasco FP-8000 fluorometer. All samples contained either 38 μM protein (wildtype) or 150 μM protein (design variant) in 25 mM sodium phosphate, 150 mM sodium chloride, pH = 7.5. Scans were acquired over a wavelength range of 400–700 nm, with excitation and emission bandwidths of 5 nm, a 50 msec response time, and a 200 nm/min scan speed.

## 8 Supplementary Figures and Tables

**Fig. S1.**
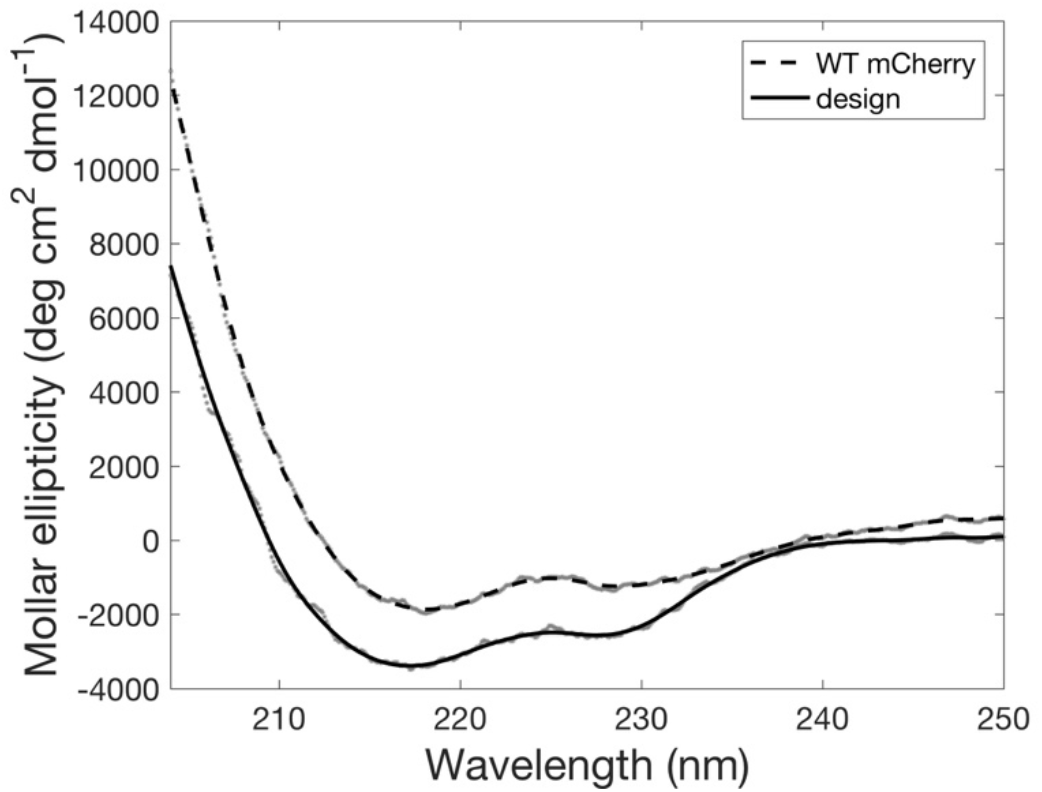
Far-UV CD spectra of wild-type mCherry and the dTERMen design. Raw data are shown with gray dots, with smooth-spline fits in dashed and solid lines for WT mCherry and the dTERMen-redesigned sequence, respectively.

**Table SI.** Native sequence recovery rates separated by core, interface, and surface positions (designated as C, I, and S, respectively).

|  | X-ray |  |  | NMR |  |  |
| --- | --- | --- | --- | --- | --- | --- |
|  | C | I | S | C | I | S |
| **dTERMen vs. native** | 36.9% | 29.7% | 24.2% | 30.3% | 24.5% | 21.3% |
| **Rosetta vs. native** | 52.1% | 30.4% | 22.9% | 33.8% | 23.3% | 17.1% |
| **dTERMen vs. Rosetta** | 31.4% | 24.5% | 25.2% | 23.9% | 20.6% | 21.7% |

**Table SII.** Structures used in native sequence recovery tests, in set I.

| **X-ray** | 3WGE | 3IDW | 1TIF | 4P0C | 1GM0 |
| --- | --- | --- | --- | --- | --- |
| 1ACF | 1COF | 3PHN | 2QMT | 3CA7 | 1QP3 |
| 1B7R | 1W42 | 4DNL | 5IJM | 3BKS | 2KRM |
| 1CHZ | 2FMA | 3OAN | 1CC7 | 2Y6X | 1KLP |
| 1F2Z | 1FX1 | 1I92 | 4GGR | 2CJ5 | 2KPK |
| 1HPT | 1TD7 | 1XAK | 2QSB | 3KT9 | 2LUS |
| 1IQ9 | 5LJK | 2FHT | 1TTZ | 4G3O | 2LQT |
| 1JWO | 2P70 | 4A60 | 1OK0 | 4HE5 | 2LSE |
| 1LN4 | 3NDQ | 3NKJ | 1NTN | 1X6J | 2KZH |
| 1P4P | 5HUA | 3LYW | 4NPE | 3LLB | 2LXY |
| 1U6T | 2BK8 | 1GUU | 4QEM | 1NB9 | 2MKW |
| 1UV0 | 3PMF | 4ICB | 1YD1 | 1KLX | 2RQS |
| 1YDL | 1A5Q | 2FU2 | 1K5A | 1DXS | 2L0C |
| 3N1R | 4GWI | 4RWU | 4EEU | **NMR** | 2LGO |
| 5FFY | 2V8L | 1WVN | 4O9F | 1AFP | 2RV8 |
| 3BJH | 2Q9X | 2IL4 | 2XL4 | 1E5C | 3GRX |
| 4NFV | 5ENB | 5I2Q | 6MSI | 1FKT | 4A4H |

**Table SIII.**
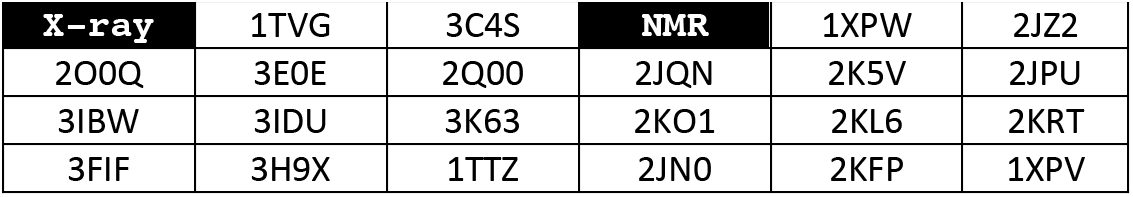
Structures used in native sequence recovery tests, in set II.

**Table SIV.**
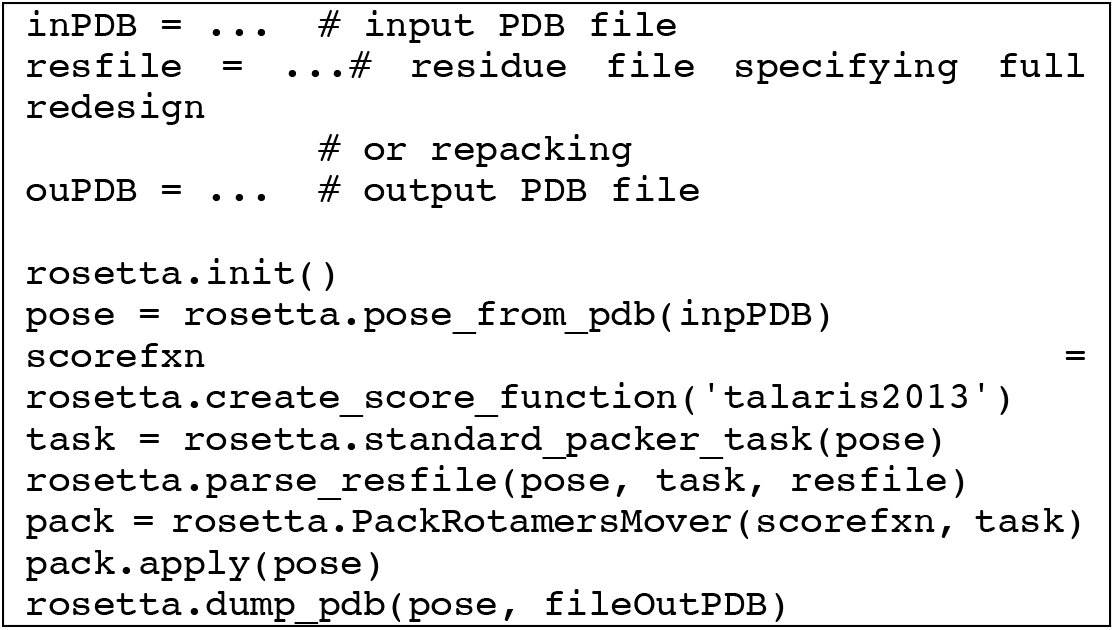
pyRosetta code used in repacking and design.

